# Mental exploration of future choices during immobility theta oscillations

**DOI:** 10.1101/2025.02.03.636313

**Authors:** Mengni Wang, Li Yuan, Stefan Leutgeb, Jill K. Leutgeb

## Abstract

Mental exploration enables flexible evaluation of potential future choices, guiding decision-making without requiring direct real-world iterations. Although the hippocampus is known to be active while imagining the future, the precise mechanisms that support mental exploration of future choices remain unclear. In the hippocampus, the theta rhythm (4-12 Hz) is prevalent during movement and supports memory coding during real-world exploration by organizing neuronal activity patterns into short virtual path segments (theta sequences) around the rat’s location. We observed these theta-related neural activity patterns during movement in a hippocampus-dependent working memory task and also, unexpectedly, theta oscillations and theta-related neural activity during immobility. Compared to standard theta sequences during movement, theta sequences during immobility differed in that they occurred at a shifted theta phase and preferentially represented remote locations, in particular the next choice in the working memory task. Coding for future locations was also observed during awake sharp wave ripple, but these short-lasting events occurred rarely and were biased toward frequently visited locations. Therefore, our findings suggest that recurring bouts of theta oscillations during immobility, which are also observed in primates and humans, support the cognitive demands of mental exploration in the hippocampal network and facilitate ongoing predictions of future choices.

## INTRODUCTION

Mental exploration enables efficient comparison of future choices and facilitates decision-making without requiring direct iterative experiences of multiple options in real life ^1–3^. During mental exploration, neural representations must dissociate from local sensory stimuli or ongoing behavior and instead be generated by internal neural circuitry to flexibly reflect remote information. Despite the importance for guiding behavior, neural mechanisms underlying mental exploration remain incompletely understood. One prominent hippocampal network oscillation during brief movement pauses—awake sharp-wave ripples (SWRs)—has been proposed as a candidate mechanism reflecting this process. The compressed neural activity patterns within SWRs can play out potential routes and predict future paths in advance of memory-guided behavior ^4–7^. However, SWR-associated neuronal firing patterns have been shown to not immediately precede memory-guided choices ^8^, and real-time disruption of awake SWRs does not impair upcoming decisions ^9^. Moreover, the sporadic nature of awake SWRs—occurring approximately once every few seconds during immobility—makes them unsuitable for supporting recursive and continuous processes required for mental exploration. This raises the question whether other types of neural activity might support more sustained coding of multiple behavioral options.

Neural coding of path segments within an environment is not limited to SWR-associated activity but has also been observed during theta oscillations, another prominent hippocampal network oscillation ^10–14^. Theta-associated coding, referred to as theta sequences, is typically confined to the vicinity of the animal’s current location, manifesting as look-ahead sweeps, flickering representations between past and future paths, or alternating left-and-right sweeps ^11, 12, 15–18^. In addition, in memory-guided tasks and/or accompanied by head movements (i.e., vicarious trial and error) ^19–21^, theta sequences have been shown to extend towards goals, represent salient locations, or alternate between potential future pathways as animals approach the branching point ^13–15, 22, 23^.

Despite these findings, the strong association between theta oscillations and movement limits their coding capabilities to active behavioral states. Although theta oscillations have also been reported during periods of high-arousal immobility—such as when behavior is driven by salient sensory stimuli and under aversive conditions ^24–29^—it remains unclear whether theta oscillations during immobility can occur independent of external stimuli and whether neural coding during theta states can be flexibly used for mental exploration across different behavioral and brain states. To investigate the mechanisms for mental exploration, we used Neuropixels probes to record hippocampal neuronal activity and oscillatory patterns in rats (*n* = 8) as they performed a hippocampus-dependent spatial working memory task ^30, 31^. Our data reveal that theta oscillations consistently emerge during movement pauses in the task in addition to periods of movement and that coding for remote trajectories is prevalent during immobility theta. These findings identify a not-yet described mechanism for generating bouts of mental exploration and reveal how theta-related hippocampal network activity facilitates memory-guided decision making.

## RESULTS

### Theta oscillations during immobility in a spatial working memory task

Although theta oscillations during immobility have been reported in numerous behavioral tasks and species, it is well-established that hippocampal theta oscillations in rodents are far more prominent during movement and that immobility often results in cessation or pauses in theta oscillations ^32, 33^. This allows for a brief window of non-theta states during and immediately after reward consumption and after stopping at other maze locations, which are accompanied by awake SWRs ^6, 34, 35^. These findings are predominantly based on recordings in or near the hippocampal CA1 layer, and therefore, we more broadly examined oscillatory patterns during immobility across all dorsal hippocampal subregions, including the dentate gyrus (DG) and CA3, by performing high-density local field potential (LFP) recordings with Neuropixels 1.0 probes (Fig. 1a, Fig. S1, *n* = 16 recording sessions in 4 male and 4 female rats). The recordings were performed in a hippocampus-dependent, and specifically, a DG-dependent spatial working memory task ^31, 36^, in which rats had to iteratively retrieve liquid reward from the 8 arms of a radial maze (Fig. 1b). In each trial, the first four arms were made available one by one in a pseudorandom order (‘forced phase’), which was followed by a phase when all arms were available and the rat had to select the four previously unvisited arms to complete a trial (‘choice phase’).

**Figure 1.**
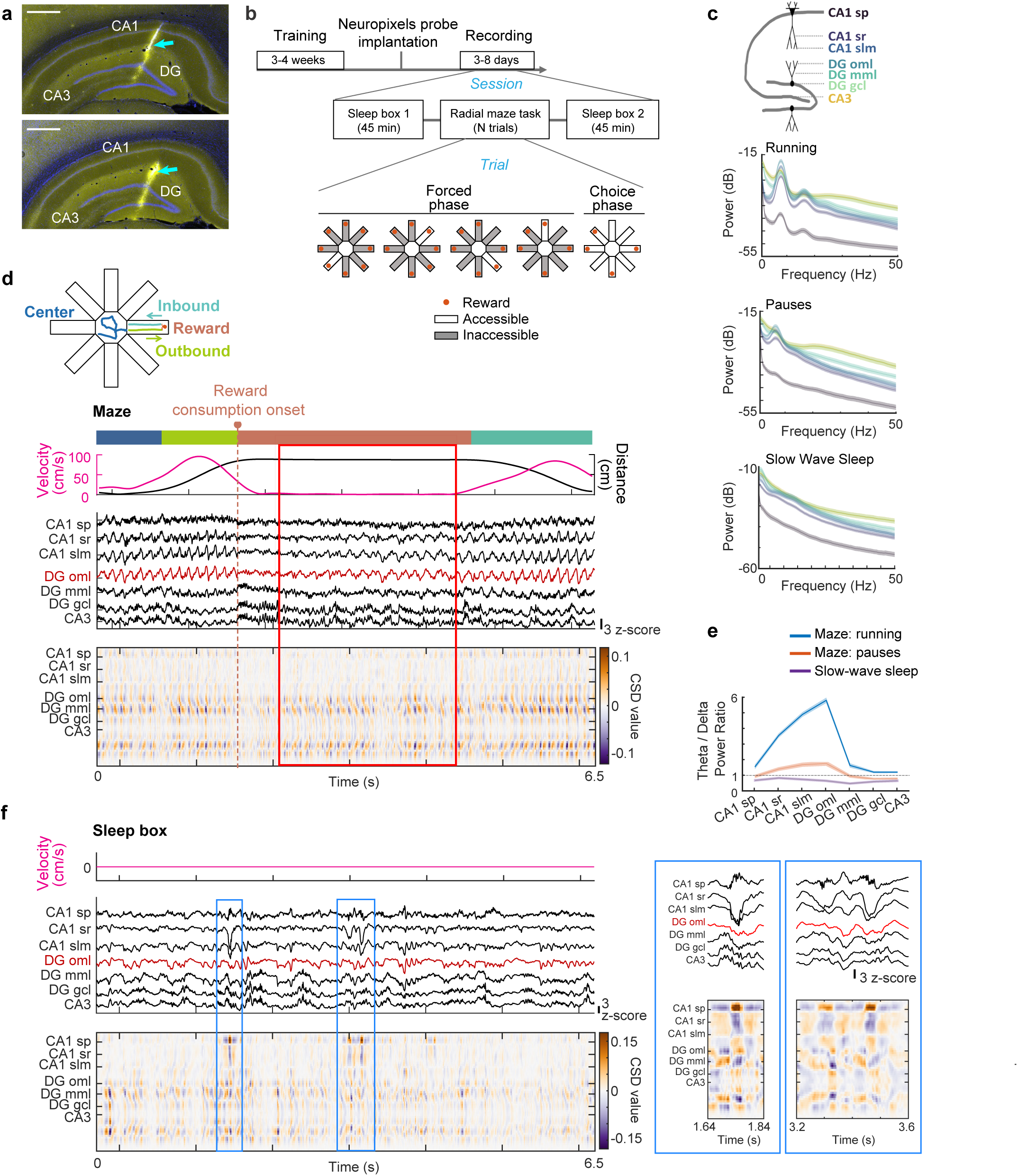
Theta oscillations in the rat hippocampus emerged during awake pauses in a memory task. **a**, Adjacent coronal brain sections through the dorsal hippocampus (HPC) of an example rat. Blue arrows highlight the Neuropixels probe track (labeled by DiI, yellow) passing through CA1, DG upper blade, CA3, and DG lower blade (labeled by DAPI, blue). Scale bar, 500 μm. **b**, Timeline of training and recording, along with a schematic of the radial-arm maze working memory task. **c**, Mean LFP spectra for seven HPC layers during maze running (velocity >10 cm/s, top), maze pauses (velocity <1 cm/s, middle), and slow-wave sleep in the sleep box (bottom). Shading represents the mean ± s.e.m. across sessions (*n* = 16 sessions for running and pauses; *n* = 14 sessions for slow-wave sleep). Theta peaks are observed in all DG layers, CA1 stratum radiatum (sr), and CA1 stratum lacunosum moleculare (slm) during pauses but are absent during slow-wave sleep. Running, theta peak frequency: 7.8 ± 0.4 Hz, mean ± sd, *n* = 16 sessions; pauses, theta peak frequency: 6.4 ± 0.3 Hz, mean ± sd, *n* = 16 sessions. Stratum pyramidale (sp), outer molecular layer (oml), middle molecular layer (mml), granule cell layer (gcl), hilar/CA3c (CA3). **d**, Behavioral and recording data from an example trial segment. Each arm visit was divided into ‘center’, ‘outbound’, ‘reward’ and ‘inbound’ epochs (with each epoch lasting from entry to exit in the four behavior zones), using the color scheme of the maze schematic. Top: Instantaneous velocity and distance to the maze center. Middle: z-scored HPC LFP traces. Bottom: LFP current source density (CSD) plot. LFP theta oscillations during immobility and the corresponding CSD patterns are highlighted with a red outline. **e**, Mean LFP theta/delta power ratio for the seven HPC layers, calculated in 2-s windows and compared between running, pauses, and slow-wave sleep. Shading represents the mean ± s.e.m. across sessions (*n* = 16 sessions, running and pauses; *n* = 14 sessions, slow-wave sleep). Theta/delta power ratios were significantly greater than 1 in all layers during running, in all layers but the CA1 sp, DG gcl, and CA3 during pauses, and in none of the layers during slow-wave sleep (*z* statistics for layers from left to right, running: 3.4, 3.5, 3.5, 3.5, 3.1, 2.3, 2.8; pauses: -1.8, 3.5, 3.5, 3.5, -1.2, -3.5, -3.5; slow-wave sleep: -3.3, -3.3, -3.3, -3.3, -3.3, -3.3, -3.3; uncorrected *P* values for layers from left to right, running: 3.5×10^-4^, 2.4×10^-4^, 2.4×10^-4^, 2.4×10^-4^, 8.8×10^-4^, 0.011, 0.0028; pauses: 0.97, 2.4×10^-4^, 2.4×10^-4^, 2.4×10^-4^, 0.89, 1.0, 1.0; slow-wave sleep: 1.0, 1.0, 1.0, 1.0, 1.0, 1.0, 1.0, one-sided Wilcoxon signed rank test). **f**, Example slow-wave sleep period in the sleep box. The two SWRs during the period are highlighted and enlarged on the right (blue boxes). Color codes and abbreviations as in d.

As expected, we observed prominent theta oscillations throughout all hippocampal subregions during task phases with running ^33, 37^, with the maximum theta amplitude below the hippocampal fissure in the dentate outer molecular layer (DG oml) (Fig. 1c and d; Running, peak theta frequency: 7.8 ± 0.4 Hz, mean ± sd, *n* = 16 sessions). During movement pauses, theta oscillations were often low or undetectable at recording sites in the CA1 cell layer. Yet, we detected prominent theta oscillations at high-amplitude recording sites (again, with a maximum in DG oml) during periods when rats were completely stopped (running speed, 0-1 cm/sec; Fig. 1c and d, Fig. S2; Pauses, peak theta frequency: 6.4 ± 0.3 Hz, mean ± sd, *n* = 16 sessions). The LFP patterns during immobility theta were clearly distinct from LFP patterns during other prolonged rest periods, which included slow-wave sleep and SWRs (Fig. 1e and f, Fig. S3, Table S1).

### The depth profile of theta oscillations across hippocampal cell layers during immobility and movement was remarkably similar

Theta oscillations during immobility have been observed in various behavioral tasks and across multiple species, and are often classified as type II theta, which is characterized by lower frequency and a higher dependence on intrahippocampal circuits for rhythm generation, compared to movement-related type I theta ^24–26, 29, 37–39^. Therefore, we asked whether any of the features of type II theta are also characteristic for immobility theta in the working memory task. First, we measured the amplitude, duration, and shape of LFP signals in different hippocampal cell layers and across velocity ranges. Theta amplitude was lower during immobility compared to movement, such that immobility theta was below or close to the detection limit in the CA1 pyramidal cell layer but not in deeper layers (Fig. 1c and e, Fig. 2b, Fig. S2). Theta cycles exhibited a shorter duration and increased asymmetry during movement compared to immobility (Fig. 2c and d; Fig. S4). The larger asymmetry was mostly a consequence of shorter descending half cycles while the ascending half showed only a minor change in duration as velocity increased (Fig. 2d).

**Figure 2.**
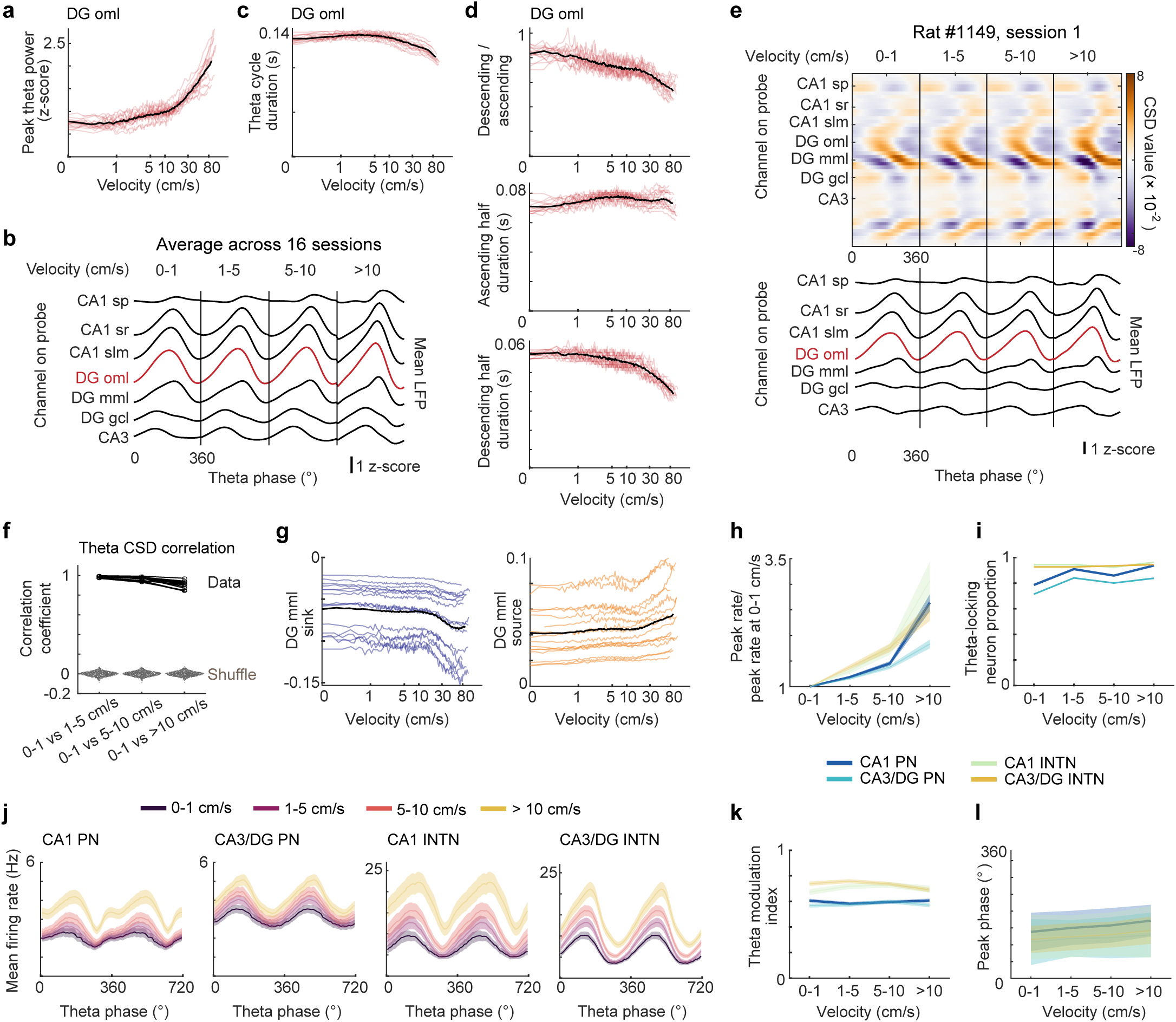
Theta oscillations during immobility compared to running were lower in frequency and amplitude, but had matching CSD patterns. **a**, Peak z-scored theta power as a function of the animal’s velocity (red lines, sessions; black line, session average, *n* = 16). Peak theta power (using the session average data) and velocity (on a log scale) were highly correlated (*r* = 0.84, *P* = 3.4×10^-27^, Pearson’s correlation). **b**, Average LFP amplitude of theta cycles (z-scored within each session and each layer, *n* = 16 sessions) for seven HPC cell layers (see Fig. 1 for layer abbreviations) and for four velocity ranges. **c**, Theta cycle duration was negatively correlated with velocity (on a log scale; *r* = -0.72, *P* = 1.8×10^-17^, Pearson’s correlation). Lines as in a. **d**, Descending-to-ascending ratio (top), duration of ascending half (middle), and duration of descending half (bottom) were negatively or positively correlated with velocity (on a log scale; *r* = -0.94, *P* =3.1×10^-47^; *r* = 0.41, *P* = 2.4×10^-5^; *r* = -0.86, *P* =2.1×10^-30^, Pearson’s correlation). Lines as in a. **e**, Top: CSD across the theta cycle, averaged over all cycles in four velocity ranges throughout an example session. Bottom: mean LFP across the theta cycle in seven HPC layers (as specified in the CSD plot). **f**, CSD patterns of theta cycles were correlated between immobility (velocity, 0-1 cm/s) and other velocity ranges (black lines, Pearson’s correlation coefficients, *n* = 16 sessions). The gray swarm charts show the shuffle distribution of correlation coefficients calculated between shuffled immobility theta CSDs (100 times, with permutations across HPC layers and theta phase) and CSDs of other velocity ranges. Correlations between data values were significantly higher than shuffled distributions (all *P* values <0.01, *n* = 16 sessions). **g**, Sink (left) and source (right) amplitudes in the DG mml layer (colored lines, sessions; black line, average over sessions, *n* = 16). The amplitudes for sinks (sources) were negatively (positively) correlated with velocity (on a log scale; *r* = -0.80, *P* = 5.1×10^-23^; *r* = 0.82, *P* = 2.0×10^-25^, Pearson’s correlation). **h**, Normalized peak firing rates (i.e., divided by peak rate at velocities <1 cm/s) of CA1 principal neurons, CA3/DG principal neurons, CA1 interneurons, and CA3/DG interneurons across velocity ranges. Shaded areas represent the mean ± s.e.m. across cells (*n* = 178, 378, 54, and 110 cells, respectively, for four cell types across all 16 sessions). The peak rates at velocities >10 cm/s were higher than at velocities <1 cm/s for all four types of neurons (*z*-statistics = 10.2, 10.8, 6.0, 8.6; *P* = 6.8×10^-25^,1.6×10^-27^, 8.8×10^-10^, 2.9×10^-18^, one-sided Wilcoxon signed rank test). Principal neurons (PN), interneurons (INTN). **i**, Proportion of significantly theta-phase locked hippocampal cells (cell types as in h) in each velocity range. Theta-phase locking of each cell was determined using the Rayleigh test for circular non-uniformity with a significance level of 0.05. The observed proportions of theta-locked cells were significantly higher than chance (CA1 principal neurons, for each of the four velocity ranges: *P* values = 9.0×10^-145^, 1.8×10^-189^, 4.9×10^-170^, 3.2×10^-201^; CA3/DG principal neurons: *P* values = 1.5×10^-257^, 0.0, 4.1×10^-316^, 0.0; CA1 interneurons: *P* values = 9.4×10^-63^, 9.4×10^-63^, 2.3×10^-60^, 2.9×10^-65^; CA3/DG interneurons: *P* values = 5.4×10^-122^, 5.4×10^-122^, 2.2×10^-124^, 7.8×10^-127^, based on binomial distribution). **j**, Mean firing rates as a function of theta phase, for each velocity range and HPC cell type (as in h). Shaded areas indicate the mean ± s.e.m. over all cells of a type. Note that each cell type exhibited similar theta-locking patterns across the velocity ranges, though firing rates increased in higher velocity ranges. **k**, Theta modulation index of a cell was defined by first calculating the average firing rate in theta phase bins and then the (maximum-minimum)/maximum across phase bins. The mean theta modulation index did not differ across velocity ranges for any cell type, except for CA3/DG interneurons (CA1 principal neurons: *F* = 0.69, *P* = 0.56; CA3/DG principal neurons: *F* = 0.97, *P* = 0.41; CA1 interneurons: *F* = 1.1, *P* = 0.34; CA3/DG interneurons: *F* = 3.5, *P* = 0.016, one-way ANOVAs). Shaded areas represent the mean ± s.e.m. across all cells of a type (as in h). **l**, The peak firing phase of all cells of a type (as in h) was averaged for each velocity range. Average peak phase of all cell types, except for CA1 interneurons, differed across velocity ranges (*F* = 10.9, 6.3, 0.45, 8.8; *P* = 5.2×10^-7^, 3.0×10^-4^, 0.71, 1.2×10^-5^, parametric Watson-Williams multi-sample test for equal means). Shaded areas represent the mean ± sd for circular data.

Next, we investigated whether neuronal activity patterns during immobility may be generated by distinct synaptic input patterns than during movement theta. To address this, we performed current-source density (CSD) analysis (Fig. 2e; Fig. S5), which is commonly used to capture synaptic input patterns in layered brain regions ^40, 41^. Despite the differences in theta amplitude and frequency, CSD analysis did not reveal distinct sink/source distribution patterns between immobility and movement theta cycles except that the magnitude of sink/source pairs in DG mml became more prominent at high velocities (Fig. 2f-g, Fig. S5). Therefore, only the magnitude, but not the distribution of inputs to hippocampus differed across velocity ranges, suggesting that immobility theta oscillations in the memory task did not share key features of type II theta, such as pronounced current sinks in CA1 ^37^. We next tested whether local interneuron and pyramidal cell populations responded differently to inputs during immobility and movement theta. While neuronal firing rates were strongly modulated by velocity (Fig. 2h,j), we found remarkably similar theta phase locking and theta phase modulation across speed ranges, with peak firing phases only slightly shifted as velocity increased (Fig. 2i-l). Along with the similar CSD patterns, this suggests that local hippocampal neurons are driven to a corresponding degree by theta-rhythmic inputs irrespective of movement velocity.

### Theta sequences persisted during immobility and preferentially represented remote locations

During active movement, hippocampal neuronal activity is thought to be organized sequentially within theta cycles, forming ‘theta sequences’ that represent short virtual spatial paths sweeping from behind the animal’s current location to tens of centimeters ahead ^10–12, 15, 16, 18, 22^. The observation of prominent theta oscillations during periods of immobility (Fig. 1), coupled with qualitative similarities in theta-associated neuronal activity between immobility and movement states (Fig. 2), suggests that sequential neuronal activity, similar to theta sequences, could extend beyond movement periods and occur during sustained periods of immobility. To test this, we analyzed the information content of dorsal hippocampal cell ensembles (*n* = 239 CA1 principal cells, 484 DG/CA3 principal cells, 62 CA1 interneurons, 136 DG/CA3 interneurons in 16 sessions from 8 rats; Fig. S6, see Table S2 for cell numbers per session) using a Bayesian probability decoding algorithm ^42–44^ that reconstructs the represented position of the rat from neural data. As expected for periods of running along tracks ^10–12, 15, 16^, we observed canonical theta sequences—decoded positions that predominantly swept ahead of the rat during each theta cycle—during inbound and outbound runs (Fig. 3a and b, Fig. S8; proportion of significant theta sequences among all qualified theta cycles during inbound/outbound run: 96.4% ± 2.0%/94.4% ± 3.3%; length of sequences during inbound/outbound run: 23.2 ± 4.0 cm/20.7 ± 3.6 cm, mean ± sd, *n* = 16 sessions).

**Figure 3.**
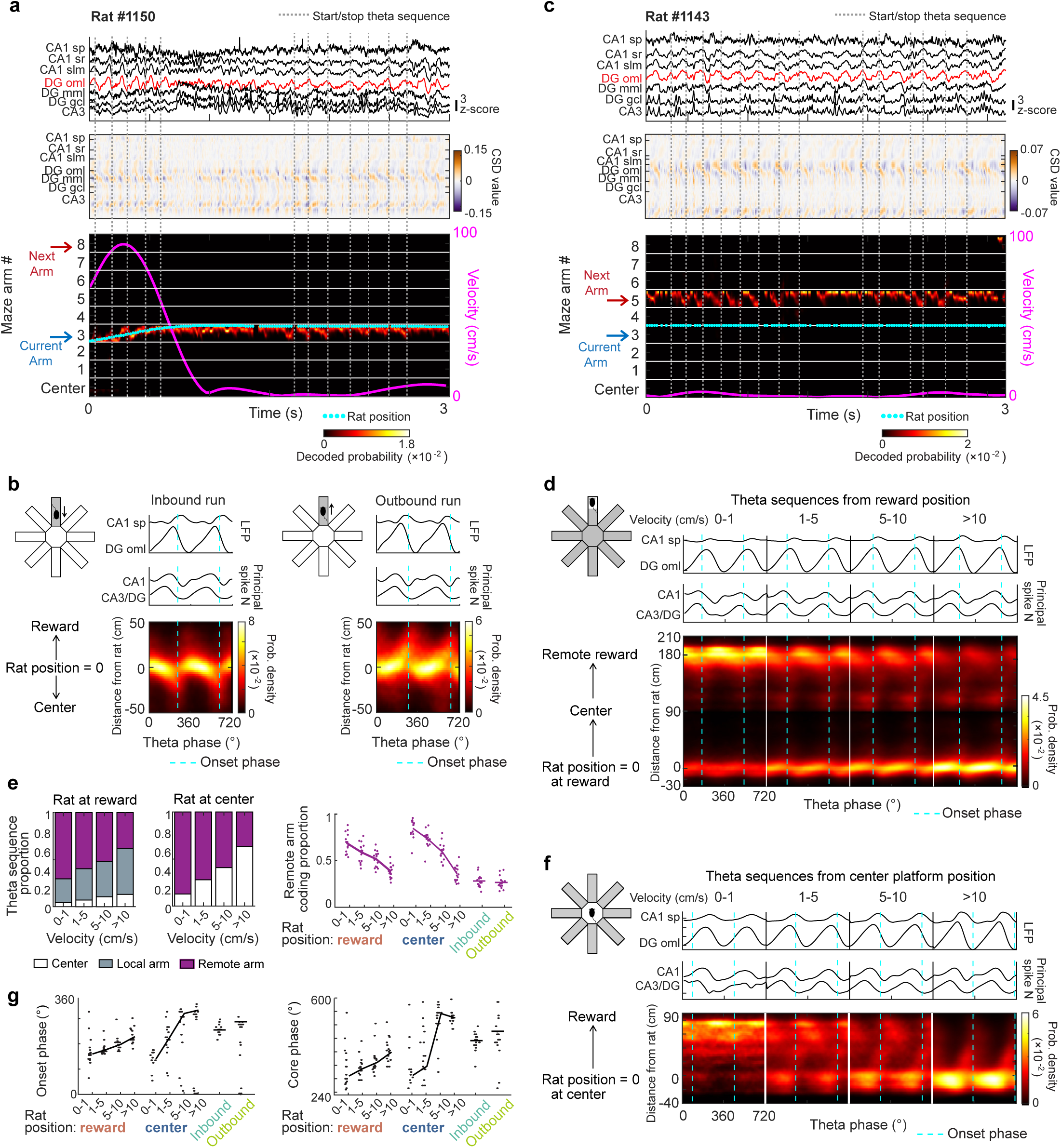
Theta sequences during immobility preferentially represented remote locations. **a**, Theta sequences (i.e., decoded spatial paths within approximately one theta cycle) around the rat’s current location were observed during outbound movement and during immobility in the reward zone. Top: Z- scored HPC LFP signals in seven dorsal HPC layers during an example trial period (red trace, DG oml; see Fig. 1 for layer abbreviations). Middle: corresponding CSD along the probe track in dorsal HPC. Bottom: Bayesian probability heatmap (20 ms decoding windows, advanced in 5 ms steps), decoded from HPC cells (see Table S2 for cell numbers and decoding accuracy, decoding error: 23.3 ± 6.8 cm, mean ± sd, *n* = 16 sessions) during the same segment as above. The y-axis represents linearized maze position in the center platform and eight arms (see Fig. S7a for the transformation from the radial maze to linear coordinates). The rat’s instantaneous velocity (pink line) and actual location (cyan dots) are overlaid. Dashed vertical lines, start/stop of clear theta sequence examples. While the onset phase of theta sequences was near the theta trough (of the DG oml LFP) during outbound runs (see theta sequences near the beginning of the example period), it was near the theta peak during immobility theta in the reward zone (see theta sequences in the second half of the example period). **b**, Standard theta sequences were observed during inbound (left) and outbound (right) runs. Top: Average z-scored LFP trace for theta cycles during the runs. Middle: Average smoothed (Gaussian filter, standard deviation = 1.2) and normalized spike counts [(spike count at each theta phase-minimum spike count within the theta cycle)/(maximum spike count-minimum spike count)] of hippocampal principal cells that were used for Bayesian decoding. Bottom: Heatmaps of locations on maze arms (relative to the rat and in the direction from the center platform to reward zones) that were maximally represented along the phases of qualified two theta cycle-long periods (*n* = 586.8 ± 72.8/416.3 ± 54.6 double cycles for inbound/outbound runs in 16 sessions, mean ± s.e.m.). The average probability density heatmap over all 16 sessions (smoothed by a 2-D Gaussian kernel, standard deviation = 2), with the heatmap showing the probability of a location being represented at a particular theta phase. Blue dashed lines, the onset phase (i.e., lowest maximal probability along the phase axis) of the average theta sequence. **c**, Theta sequences at remote locations were observed during immobility theta. In the example trial segment (see a for labels and organization), the representation pointed to an unvisited remote arm while the rat was stopped in the current reward zone. The represented remote arm corresponded to the next arm in the sequence of arm visits. Note the onset phase of theta sequences near the theta peak during immobility theta in the reward zone. **d**, With the rat in the reward zone, remote locations were preferentially represented during immobility theta, while the current location was gradually more represented as velocity increased (*n* = 1567.4 ± 173.1, 3958.1 ± 234.5, 2365.0 ± 139.7, 2457.5 ± 171.7 qualifying double cycles in four velocity ranges, mean ± s.e.m., *n* = 16 sessions). Panel layout as in b. **e**, Proportion of theta cycles that predominantly represented the center platform, current arm, or remote arms. Data are shown separately for each velocity range and with the rat either in the reward zone (left) or at the center platform (middle; reward locations: *n* = 51740, 106645, 62072, and 69187; center platform: *n* = 6390, 15600, 14085, and 70410 qualifying theta cycles in four velocity ranges, *n* = 16 sessions). Right: Proportions of theta cycles with preferential coding for remote arms are plotted by sessions (dots), when rats were in the reward zone or on the center platform (as in the panels to the left) and also when the rats were running outbound or inbound (lines, average over sessions within each zone/velocity range). Running on maze arms is mostly at velocities >10 cm/s, and therefore, lower speed ranges are not shown. With the rat at reward or center, the proportion of remote coding is negatively correlated with running velocity (Reward: *r* = -0.78, *P* =1.9×10^-14^; Center: *r* = -0.89, *P* = 1.6×10^-22^; Pearson’s correlation). **f**, With the rat in the center, remote locations were preferentially represented during immobility theta and current locations were increasingly represented at higher velocities (*n* = 124.9 ± 46.1, 388.2 ± 74.9, 393.6 ± 60.4, 2299.7 ± 130.4 qualifying double cycles in four velocity ranges, mean ± s.e.m., *n* = 16 sessions). Panel layout as in b. **g**, Onset phase (left) and core phase (right) of theta sequences as a function of the rat’s velocity and behavior phase. Sessions, dots; lines, circular mean over sessions within each zone/velocity range. With the rat at reward or center, onset and core phases were correlated with the rat’s velocity (onset phase, reward: rho = 0.70, *P* = 1.3×10^-7^; center: rho = 0.81, *P* = 6.1×10^-9^; core phase, reward: rho = 0.57, *P* = 3.2×10^-5^; center: rho = 0.70, *P* = 5.5×10^-7^; circular-linear correlation).

In addition to the standard theta sweeps during running, we asked whether similar sequential neural representations could also be decoded during immobility theta. We first focused on pauses in the reward zones and observed clear spatial sweeps during immobility theta, similar to theta sequences during movement (Fig. 3a, Video S1). Interestingly, neural coding during immobility theta was not restricted to the proximity of the rat’s current position (’local’ theta sequence), but frequently covered a different maze arm than the one the animal was currently occupying (‘remote’ theta sequence) (Fig. 3c- e, Fig. S7, Fig. S9, Video S2; proportion of significant local/remote theta sequences at reward when velocity <1 cm/s: 29.4% ± 10.7%/64.4% ± 11.4%; length: 14.2 ± 2.7 cm/17.8 ± 4.4 cm, mean ± sd, *n* = 16 sessions). We next analyzed neural coding on the center platform and observed prominent theta sequences, again, with a preference for representing remote locations during immobility (Fig. 3e and f, Fig. S9a-h; proportion of significant local/remote theta sequences when velocity <1 cm/s: 11.9% ± 12.5%/85.1% ± 14.0%; length: 28.2 ± 13.8 cm/18.7 ± 4.2 cm, mean ± sd, *n* = 16 sessions). Given that theta sequences frequently represented remote locations during immobility while typically representing proximal locations during running, we sought to determine to what extent the movement state controlled the relative balance of local compared to remote representations. We found that remote representations by theta sequences became less frequent as the animal’s instantaneous velocity increased (Fig. 3e; Pearson’s correlation between velocity and remote-arm coding proportion, Reward: *r* = - 0.78, *P* = 1.9×10^-14^; Center: *r* = -0.89, *P* = 1.6×10^-22^). Interestingly, theta sequences at lower velocities predominantly occurred at a shifted phase range compared to theta sequences at higher velocities, including during inbound and outbound movement on the track (Fig. 3g, Fig. S9e,f; circular-linear correlation between onset phase and velocity, reward: rho = 0.70, *P* = 1.3×10^-7^; center: rho = 0.81, *P* = 6.1×10^-9^; onset phase of theta sequences at reward/center when velocity <1cm/s: 144.3 ± 47.0°/135.9 ± 37.4°, during inbound/outbound run: 253.6 ± 19.7°/286.0 ± 63.6°, mean ± sd, *n* = 16 sessions). Taken together, our findings demonstrate that hippocampal neuronal activity was organized into theta sequences during immobility theta, similar to the sequential neural representation during standard movement theta sequences. Yet, immobility theta sequences differed from their movement counterpart in that they occurred at a shifted theta phase range and preferentially coded for distant remote locations rather than the vicinity of the rat’s current location.

### Theta sequences during immobility at reward sites and choice points predicted the upcoming choice

The flexible and pronounced representation of remote arms during prolonged pauses at reward sites and choice points (Fig. 3c-f) raised the question whether theta sequences during immobility contribute to planning and predict the animal’s future choices in the task. To investigate this, we tested whether the next arm was represented more frequently than other remote arms during the choice phase (Fig. S10). During theta states on the center platform, the upcoming arm choice was significantly overrepresented immediately before the animal moved onto that arm (Fig. 4a, right), consistent with previous studies showing that future paths towards a goal were represented in theta sequences when approaching choice points of tasks ^13–15, 22^. Interestingly, predictive neural representations for the next reward could be traced back to the preceding reward location, which is positioned ∼180 cm away from the next reward location. The predictive code at the preceding reward emerged at lower coding probability than during subsequent theta bouts on the center platform, when it increased immediately preceding the next arm entry (Fig. 4a, left, Fig. S7; Video S2). Notably, the predictive representation of the next arm in the reward zone and at the subsequent choice point was positively correlated (Fig. 4b), suggesting a progression of coding strength for an upcoming arm choice during decision-making. The preferential representation of the next choice was absent during the forced phase, when the animal followed the available path without making a choice.

**Figure 4.**
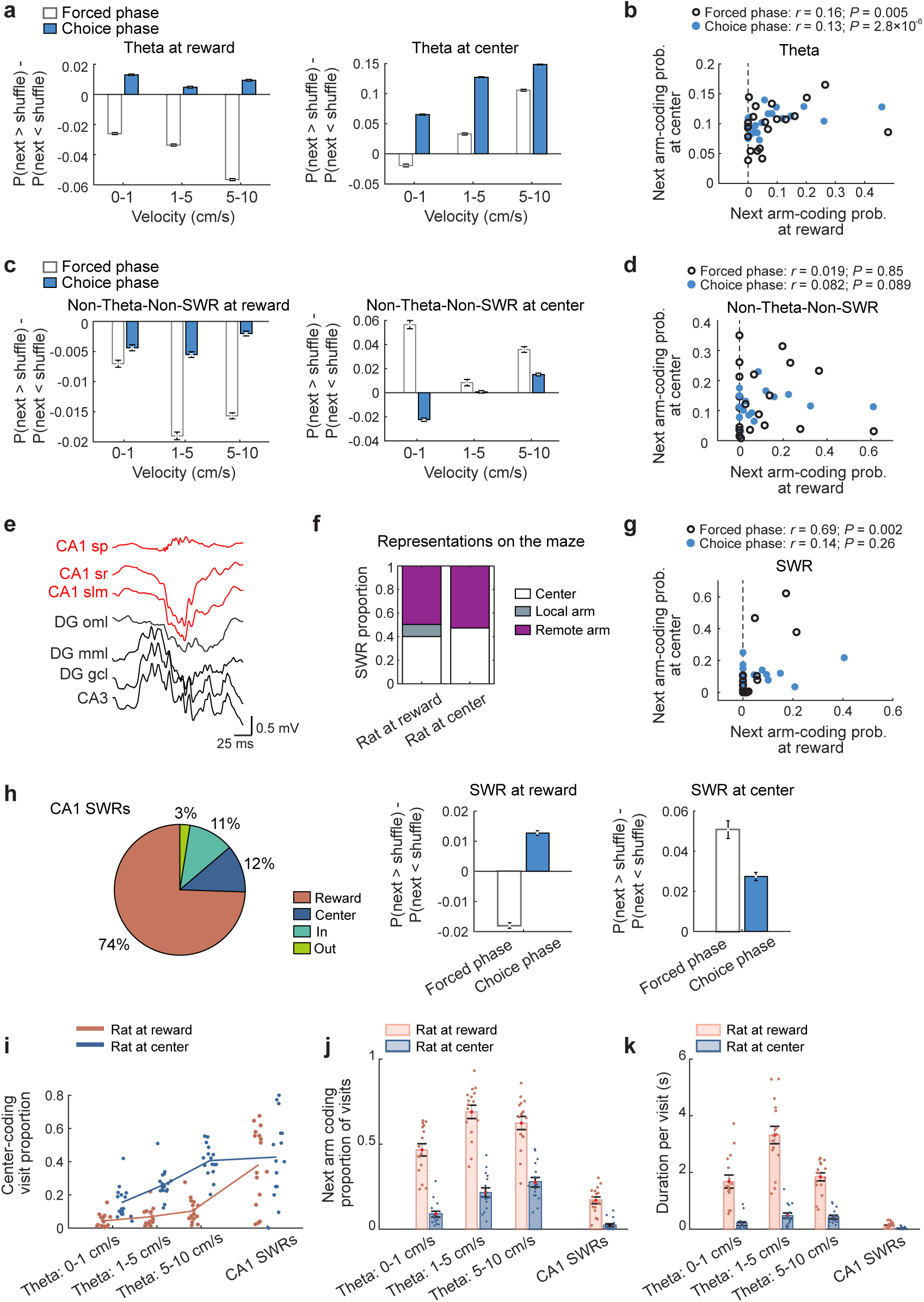
Next choice predictions during immobility theta were more sustained than during CA1 SWRs. **a**, When rats were in the reward zone, the coding probability for the next arm was higher than for shuffled arms (*n* = 500 shuffles, includes all arms other than the current and next one, method details in Fig. S10) during the choice, but not forced phase (left; forced: *t* = -40.45, -45.38, -77.83; *P* = 1.0, 1.0, 1.0; choice: *t* = 25.07, 8.59, 16.85; *P* = 1.1×10^-90^, 5.5×10^-17^, 4.8×10^-51^ for each of the three velocity ranges, one-sided *t*-test for next > shuffles). When at the center, coding for the next arm was higher than for shuffled arms during the forced and choice phase (right; forced: *t* = -8.36, 19.24, 66.62; *P* = 1.0, 2.0×10^- 62^, 8.3×10^-251^; choice: *t* = 62.52, 148.0, 187.8; *P* = 1.7×10^-238^, 0.0, 0.0, one-sided *t*-test for next > shuffles). **b**, Probabilities of next arm representations were correlated between pairs of epochs in the reward zone (at velocities <10 cm/s) and at the subsequent pass through the center platform (at velocities <10 cm/s; see figure for statistics, Spearman’s rho, *n =* 294/1288 reward-center pairs with sufficient decoding sampling at both sites in forced/choice phase). For visualization, raw data was grouped into 20 groups based on the next arm representation probability in the reward zone (x-axis), and the mean probability for next arm coding in the center was plotted for each group (y-axis). **c**, Calculated as in a, but for ‘Non-Theta-Non-SWR’ periods, which were defined as periods with z-scored theta power <-1, CA1 SWR events excluded, and velocities <10 cm/s. Coding for the next arm was higher than coding for shuffled arms when rats were on the center platform, but not when at reward locations (reward, forced: *t* = -12.00, -29.72, -31.48; *P* = 1.0, 1.0, 1.0; choice: *t* = -8.71, -11.64, -5.00; *P* = 1.0, 1.0, 1.0; center, forced: *t* = 16.26, 3.10, 15.02; *P* = 2.6×10^-48^, 0.001, 1.3×10^-42^; for choice: *t* = -15.48, 0.62, 13.34; *P* = 1.0, 0.27, 3.1×10^-35^ for each of the three velocity ranges, one-sided *t*-test for next > shuffles). **d**, Calculated as in b, but for ‘Non-Theta-Non-SWR’ periods. Probabilities of next arm representations were not correlated between pairs of epochs in the reward zone and at the subsequent pass through the center platform (see figure for statistics, Spearman’s rho, *n* = 98/427 reward-center pairs with sufficient decoding sampling at both sites in forced/choice phase). **e**, LFP traces across seven HPC layers during an example CA1 SWR. Red lines indicated channels used for identifying SWR events. **f**, Proportion of ripples during which the center platform, current arm, or remote arms were preferentially represented while animals were in the reward zone or at the center platform (reward/center: *n* = 6465/1094 CA1 SWRs across 16 sessions). **g**, Calculated as in b, but for SWRs. In the choice phase, probabilities of next arm representations were not correlated between pairs of epochs in the reward zone and at the subsequent pass through the center platform (see figure for statistics, Spearman’s rho, *n* = 17/64 reward-center pairs with sufficient decoding sampling at both sites in the forced/choice phase). **h**, Left: percentage of CA1 SWRs in four behavioral zones. Middle and right: Calculated as in a, but for SWRs. Coding for the next arm was higher than coding for shuffled arms except for the reward zone/forced phase combination (reward, forced and choice phases: *t* = -19.87, 16.45; *P* = 1.0, 3.5×10^-49^; center, forced and choice phases: *t* = 11.49; 13.35; *P* = 1.4×10^-27^, 2.9×10^-35^, *n* = 16 sessions, one-sided *t*-test for next > shuffles). **i**, Proportion of theta cycles and SWRs that predominantly represented the center platform are plotted by session, and for theta cycles, by velocity range (colored dots). The center coding proportion was higher in SWRs than in theta cycles, except for the 5-10 cm/s velocity/rat at center combination (reward: *z* = 3.9, 3.5, 3.1; *P* = 4.8×10^-5^, 2.1×10^-4^, 0.0011; center: *z* = 3.2, 2.1, 0.26; *P* = 5.9×10^-4^, 0.019, 0.40 for the three velocity ranges; *n* = 16 sessions, one-sided Wilcoxon rank sum test). Colored lines, mean over sessions within each category. **j**, Proportion of behavior epochs when the next arm was represented. The next-arm coding proportion was higher in theta cycles than in SWRs with the rat in the reward zone or at the center platform (reward: *z* = 4.5, 4.8, 4.7; *P* = 3.3×10^-6^, 7.7×10^-7^, 1.1×10^-6^; center: *z* = 3.5, 4.6, 4.8; *P* = 2.4×10^-4^, 1.9×10^-6^, 8.9×10^-7^ for the three velocity ranges; *n* = 16 sessions, one-sided Wilcoxon rank sum test). **k**, Cumulative duration of theta cycles and SWRs per behavioral epoch. The duration of theta states was longer than the duration of SWR states when the rat was in the reward zone or at the center platform (reward: *z* = 4.8, 4.8, 4.8; *P* = 7.7×10^-7^, 7.7×10^-7^, 7.7×10^-7^; center: *z* = 3.5, 4.6, 4.7; *P* = 2.1×10^-4^, 1.9×10^-6^, 1.6×10^-6^ for the three velocity ranges; *n* = 16 sessions, one-sided Wilcoxon rank sum test).

To determine whether the next-choice prediction at the reward zone and center platform was specific to theta states or occurred across all brain states, we performed a similar analysis during Non-Theta-Non-SWR periods, characterized by low LFP theta power and the absence of CA1 SWRs during immobility. During these periods, we did not observe significant representation of the next arm in the reward zone or a correlation between next-choice prediction at reward and choice points (Fig. 4c,d). In addition, we analyzed retrospective coding for the previous arm and found that the previous arm was not preferentially represented during any brain state (i.e., Theta, SWRs, and Non-Theta-Non-SWRs; Fig. S11). These findings suggest that immobility theta sequences at reward reflected the early planning and prediction of the animal’s future choices and were informative well in advance of reaching choice points and future goals in the task.

### Network activity patterns during CA1 SWRs and immobility theta were distinct

Awake SWRs are known to predominantly occur around the time of reward consumption, and during awake SWRs, synchronous hippocampal population activity represents remote locations ^4–7^, as was the case for population coding during immobility theta. The manifestation of remote coding during both types of oscillations raises the question whether theta and sharp-wave ripples co-occurred or whether these brain states were mutually exclusive. CA1 SWRs occurred when theta was at its lowest power, and periods of immobility theta without SWRs had a depth profile that was very distinct from SWRs (Fig. S12). In instances when CA1 SWRs were identified along with higher theta power, the CSD profile of SWRs was analyzed and found to be highly similar to those of CA1 SWRs during non-theta states in the maze and during rest periods (Fig. S12c). Therefore, coincident detection of the two network patterns is an artifact of the sharp wave generating power in the theta band rather than of SWRs occurring during sustained theta periods. To be certain that SWRs with theta power generated by sharp waves were not included as false positives in our theta analysis, we had already excluded all events with ripple power from theta periods in any of our preceding analyses. Conversely, when analyzing SWRs, we confirmed that SWRs occurred predominantly at reward sites in the spatial working memory task, coded for remote locations, and predicted future choices of the animal (Fig. 4e-h), as previously reported ^4–7^.

### Remote coding was more prominent during theta oscillations than during SWRs

Given that the prominent prediction of the animal’s upcoming choices was observed during two distinct and incompatible network states—theta oscillations and SWRs—we sought to investigate the distinctions between these two coding schemes. First, while both theta oscillations and SWRs significantly predicted the animal’s next choice (Fig. 4a, h), the systematic buildup of predictive coding capability observed for immobility theta between the reward site and the subsequent choice point (Fig. 4b) was not evident for SWRs (Fig. 4g). This discrepancy likely reflects the predominance of SWRs at reward sites along with their infrequent occurrence at choice points, precluding a mechanism for progressive predictive buildup. Second, the two oscillatory states exhibited distinct spatial coding preferences. At reward sites, neural activity during immobility theta predominantly coded for arm locations with minimal representation of the center platform, whereas SWRs showed a marked representing preference for the central platform, which was conceptually the ‘base’ location in the task and thus most frequently visited (Fig. 4f and i, Fig. 3e). Third, SWRs in CA1 occurred only during a small fraction of visits to reward locations and the center platform, resulting in inconsistent brief temporal windows for coding future locations compared to the much more prevalent theta cycles (Fig. 4j and k). The recurrent and continuous nature of theta oscillations, therefore, enables a more robust and iterative mechanism for representing upcoming choices, consistent with extended periods of mental exploration.

## DISCUSSION

Imagining the future and mental exploration are cognitive functions that are supported by episodic memory systems, including the hippocampus ^1–3, 45, 46^. Here, we observed that hippocampal theta oscillations, which support memory coding during active exploration ^15, 21, 37, 47, 48^, extended beyond movement and persisted during immobility periods. During immobility theta, hippocampal neuronal activity was organized into theta sequences, similar to the sequential neural representation during standard movement theta sequences ^11, 12, 22^. Yet, immobility theta sequences differed from their movement counterpart in that they occurred at a shifted theta phase range and preferentially coded for distant remote locations. Because immobility theta sequences were not as tightly bound to representing locations in the vicinity of the rat’s current position, they could recursively play out multiple arm choices in the task, which is reminiscent of a virtual exploration of possible future choices. Accordingly, decoded locations during immobility theta preferentially pointed to the next choice in the sequence of visited locations. The preferential coding for the next arm emerged at the previous reward location and was then retained and augmented during the subsequent pass through the center platform, immediately preceding the arm choice. As previously reported ^4–7^ we also found reactivation of remote locations and predictive coding of future choices during awake SWRs in the memory task. However, SWRs occurred only during a small fraction of visits and lasted for shorter durations than immobility theta. Therefore, the higher incidence and extended duration of theta periods in the hippocampal network make them better suited for recursive and sustained predictions of future choices and for supporting the cognitive demands of mental exploration.

Theta oscillations are the most prominent oscillatory brain activity during movement in rodents, while other species, including humans and non-human primates, show only brief bouts of theta oscillations that are not strongly tied to locomotion ^49–54^. Here we found that, even in rodents, theta is not consistently tied to movement in a memory task. The association of theta oscillations with immobility could be seen as consistent with a different type of theta—type II—which has been reported to emerge with sensory stimulation during immobility and is atropine sensitive ^24, 25, 37, 38^. Compared to movement—or type I—theta ^33, 37^, type II theta is not only characterized by its association with immobility, but also by a lower frequency range, a similarly pronounced amplitude in CA1, more marked CSD sources in CA1, and possibly, a more marked generation in ventral hippocampus ^24, 37, 38, 55, 56^. While the immobility theta in the memory task shares one of these features of type II theta—the lower oscillation frequency compared to type I theta—there are also substantial differences. The theta amplitude in the CA1 pyramidal cell layer was low, such that immobility theta was readily detectible only in other hippocampal layers, in particular those where entorhinal inputs terminate (i.e., stratum lacunosum moleculare and the dentate molecular layers). Accordingly, CSD analysis of immobility theta identified prominent sinks in these layers, which is distinct from type II theta, which is thought to be associated with pronounced sinks in CA1 layers ^37, 38^. Furthermore, the similar input distribution between memory-related immobility theta and movement theta makes these types of theta distinct from the input patterns reported for respiratory-related oscillations, which can also be detected in hippocampus ^57^. Therefore, our findings do not merely confirm that theta in rodents can occur during immobility, but also suggest that immobility theta cannot be categorized as a single theta type. For example, sensory-evoked immobility theta requires the presentation of sudden and/or arousing sensory stimuli ^24, 25^, whereas memory-related immobility theta seems to be internally generated without unexpected sensory stimuli. While cataloguing the various types of oscillations in the theta range will require a full characterization in each behavioral task, the memory-related theta oscillations that we describe here have the distinct identifying features that they are consistent with medial entorhinal inputs to the dentate middle molecular layer and are associated with remote coding for future goal locations. The importance of entorhinal inputs to DG in predicting future goals is consistent with the recently identified role of dentate circuits in prospective coding ^31, 58^. Furthermore, given that theta oscillations in humans and primates are intermittent, that theta oscillations increase during deliberate decisions and that the human hippocampus is critical for imagining the future ^1, 2, 21, 45, 46, 52–54, 59^, our related findings in rodents suggest that oscillations in the theta frequency range are associated with mental exploration across species.

The occurrence of theta oscillations during immobility raises the question whether theta states might overlap with awake SWRs, which are also selectively generated during movement pauses ^6, 34, 35^ and thus in the same behavioral state as immobility theta. Despite the association of each of these oscillation patterns with immobility, immobility theta and SWRs were mutually exclusive. SWRs were restricted to non-theta periods and displayed a pattern of synaptic inputs (e.g., large sink in stratum radiatum) that was clearly distinct from theta oscillations (see Fig. 1f, Fig. 2e, Fig. S12). While the two types of oscillations were separate in time, they shared the feature that future or past locations could be represented ^4–7, 10–16, 18^. The hippocampal population activities during SWRs and during theta states, therefore, include representations of maze locations far from the animal’s current spatial position, such as to distant reward arms. The finding that remote coding can occur during either SWRs or theta oscillations raises the question whether the two types of oscillations could interchangeably guide decision making and play out routes to upcoming choices in memory tasks ^4, 6, 7^. For example, neural activity patterns during awake SWRs can code for a future path in advance of the execution of memory-guided behavior ^5^. However, replayed sequences in SWRs do not only include immediately upcoming journeys, but also those further in the future or towards familiar locations. Furthermore, it has recently been shown that immediately upcoming behavioral choices are not preferentially activated during awake SWRs ^8^ and that real-time disruption of awake SWRs does not impair immediately upcoming memory-guided behavior ^9^. Rather than coding for upcoming choices, the key role of awake SWRs, in particular at reward locations ^60^, could therefore be an initial strengthening and updating of sequences, which then continues in subsequent rest and sleep periods ^61–65^. In support of a critical role of awake SWRs in supporting the stabilization of memory over longer intervals, it has been shown that selective elimination of awake SWRs in hippocampus impairs consolidation ^66^ and slows acquisition of a spatial working memory task ^7^ while prolonged SWRs are associated with improved retention in memory tasks^67^. Reactivation and replay during awake SWRs can therefore be seen as the initial step towards further stabilization and consolidation in subsequent sleep events ^7, 61–65^ rather than as guiding upcoming choices.

In contrast to the role of SWRs in stabilizing memory content over longer periods, remote coding during immobility theta began to show a preference for the upcoming choice following the consumption of the previous reward. The activity then re-emerged immediately before the next choice and is thus well positioned to support upcoming memory-based decisions. While SWRs may therefore have a key function for memory consolidation and for broadcasting hippocampal information to the cortex ^7, 61–65^, our findings indicate that theta sequences during immobility may serve as the primary mode for supporting memory-based decisions during ongoing behavior while using information that is received from entorhinal inputs to the hippocampus. Similar coding for future choices was not observed during non-theta periods, consistent with the well-established role of theta oscillations before and during the choice phase of spatial working memory tasks, when disruption of theta by medial septal manipulations is particularly deleterious ^48, 68^. Accordingly, theta oscillations are fundamental for memory computations, and in addition to the proposed switching between encoding and retrieval within theta cycles during movement ^47^, the continuous nature of immobility theta in memory tasks allows for an iterative mechanism for representing upcoming choices and supporting extended periods of mental exploration.

## ACKNOWLEDGEMENTS

We thank Mia Anderson, Priscilla Ee, Junhao Zhu, and Aneesh Swamy for technical assistance. This work was supported by NIH grants MH119179 to J.K.L., NS084324, NS102915, NS121231 to S.L., NSF grant 2024776 to S.L., and a Kavli Institute for Brain and Mind Postdoctoral Fellowship to M.W.

## AUTHOR CONTRIBUTIONS

J.K.L., S.L. and M.W. designed the experiments, conceptualized analyses, interpreted data and wrote the manuscript. M.W. acquired data, designed analyses, wrote and curated analysis code, and analyzed the data. L.Y. developed the recording methods. J.K.L and S.L supervised the project.

## CODE AVAILABILITY

Custom code for processing the data reported in this paper will be available from Github. Standard code that is not novel to this study will also be available from the corresponding authors upon request.

## DATA AVAILABILITY

Source data will be provided with this paper. Additional data that support the findings of this study are not in standardized formats, but can be made available without restrictions upon request to the corresponding authors.

## METHODS

### Approvals

All surgical and experimental procedures were conducted at the University of California, San Diego, as approved by the Institutional Animal Care and Use Committee and according to National Institutes of Health guidelines.

### Subjects

Female (*n* = 4, 7 weeks old) and male (*n* = 4, 9 weeks old) Long-Evans rats were obtained from Charles River Laboratory. The animals were housed individually and maintained on a 12-hour reversed light-dark cycle with lights off at 7:00 a.m. Behavioral training and in vivo electrophysiological recordings were performed in the dark phase. Following at least 1 week of adaptation to the laboratory, access to food was restricted and the rats were maintained at ∼85% of free-feeding body weight. Water was readily available. Animals were habituated and trained in a spatial working memory task ^31^, one session per day, for 3-4 weeks prior to surgery. At the time of surgery, animals were 3-5 months old (weight, females: 234 ±10 g, males: 380 ±41 g, mean ± sd). Electrophysiological recordings in the behavioral task were performed 4-5 days after surgery.

### Behavioral task

Rats were trained to perform a DG-dependent spatial working memory task on an eight-arm radial maze ^30, 31^ (Fig. 1b). A semiautomated black Plexiglas maze consisted of a central circular platform (27 cm diameter) and eight connecting arms (79 cm × 12 cm) to which access was controlled by a remote switch. A reward (chocolate milk, 0.2 mL) was available at the end of each arm. First, daily 10-min habituation sessions with immediate access to all eight arms were performed until the rat learned to consume the reward at the end of each arm. In the next training stage, each trial started with access to all arms initially restricted. The first arm was made available by the experimenter, followed by a sequence of 3 additional arms presented one at a time (forced phase). The sequences during the forced phase were pseudorandom, excluding repeated arms and instances in which all four arms would have been presented in a sequential order (e.g., 1, 2, 3, 4). Immediately after the forced phase, all eight arms were presented simultaneously (choice phase). During the choice phase, the animal was expected to visit the remaining four baited arms, and reentries into previously visited arms were considered errors. A trial ended when the rat had retrieved the reward from all eight arms and returned to the center platform or when the total time exceeded 10 min. Trials that exceeded the time limit were removed from analysis. After an intertrial interval of 2 min, the next trial was initiated. Training time was gradually increased in line with the animal’s improved performance over a 2-3 week training period, starting from 30 min up to 1 hour per day. Well-trained animals were defined as those completing at least 20 trials within a 1-hour period with an accuracy rate above 80%. Once the animals reached this criterion, we performed an additional week of training with two modifications: (1) A sleep session was added before task training to habituate the animals to the sleep box (constructed from transparent plastic, 30 cm by 30 cm; height, 50 cm) used in the subsequent recording experiments. (2) The training duration was extended to 1.5–2 hours to ensure stable task performance. Trained animals were provided free access to food for at least 3 days prior to surgery.

### Surgical procedures

Surgeries was performed using sterile procedures. Anesthesia was maintained throughout surgery with isoflurane gas (0.8%-2.0% isoflurane delivered in oxygen at 1 L/min) and with body temperature and breathing rate periodically monitored. Buprenorphine (0.02 mg/kg) was administered as an analgesic. Animals were positioned in a Kopf stereotaxic instrument, and incisor bars were adjusted until bregma was level with lambda. Implantation coordinates were developed to target 3 subregions of the hippocampus (CA1, CA3, dentate gyrus) in dorsal and ventral hippocampus with a single 1.0 Neuropixels probe (ML: -1.3 mm, AP: -3 mm for male rats; ML: -1.5 mm, AP: -3 mm for female rats, angled by 32.5° lateral from the ML axis in the ML-AP plane and 30° posterior from the DV axis in the DV-AP plane). Probes were mounted on a drive (3Dneuro) and the probe shank was dipped in 2% DiI solution (ThermoFisher) at least 5 minutes before implantation to enable tracking of the probe’s position in the brain later with histology. The shank was slowly lowered for up to 9.3 mm into the brain, the craniotomy was filled with sterile wax, and dental cement secured the drive to skull screws and sealed the incision. One of the screws was used as a ground screw and wired to the grounding and reference pads of the Neuropixels probe. The probe and the probe holder were protected by a 3D printed shroud ^69, 70^. All animals received postoperative care for at least 5 days after surgery, followed by daily health checks until the animals were euthanized. After completing all recordings for a rat, the Neuropixels probe was carefully extracted under isoflurane anesthesia before injecting an overdose of pentobarbital and performing a transcardial perfusion in a deeply anesthetized state, first with saline and then with 4% paraformaldehyde. The brain was extracted, fixed in 4% paraformaldehyde for at least 24 hours and subsequently transferred to a 30% sucrose solution until it sank, indicating full cryoprotection. Once fully cryoprotected, the brain was frozen with dry ice and sectioned coronally at a thickness of 40 μm along the entire extent of the hippocampus. DAPI staining was applied to the mounted brain slices, which were imaged using an Olympus VS200 slide scanner equipped with the Olympus VS-ASW software to visualize DAPI and DiI fluorescence images. The probe position was reconstructed in post-mortem histological material by marking the DiI fluorescence in the serially sectioned brain slices, as detailed in Fig. S1. However, estimates of anatomical locations were subject to a degree of measurement error due to the limitations of the alignment process. Therefore, the final detailed location of hippocampal layers along the probe was determined by LFP and spike analysis, as described below (see Dorsal hippocampus layer estimation).

### Chronic electrophysiological recordings in awake-behaving rats with Neuropixels 1.0 probes

After implanted animals recovered from surgery, behavioral experiments resumed. Once the animals performed >20 trials within 1.5-2 hours, electrophysiological recording sessions were performed for a minimum of 2 days. Behavioral procedures in the working memory task were identical to those without recordings except that each recording session began and ended with a 45 min rest period in the sleep box flanking the spatial working memory task. Using a video camera connected to a Digital Lynx data acquisition recording system (Neuralynx, USA), animal position was monitored by tracking an LED (sampling rate 30 Hz) that was attached to the implant. The data acquired with the Digital Lynx and Neuropixels 1.0 recording systems were synchronized in time by a 0.5 Hz TTL pulse from an Arduino. The Neuropixels 1.0 implant was connected to the interface with a tether that was counterbalanced by an elastic to absorb torque and movement during behavior on the maze. Neuropixels 1.0 probes allowed for the continuous sampling from 384 channels in two formats: action potential (AP) data (digitized at 30 kHz with gain of 500, 250, or 125, adjusted for each animal to maintain data saturation below 1%) and local field potential (LFP) data (digitized at 2.5 kHz with gain of 250 or 125, adjusted for each animal to maintain data saturation below 1%). SpikeGLX software (https://billkarsh.github.io/SpikeGLX/) was used to control acquisition and configure the probes. The positioning of an implanted Neuropixels 1.0 probe across hippocampal layers was estimated in live animals during a test recording session in the sleep box. In this session, recording channels were selected to be spatially evenly distributed along the probe. The location of the CA1, DG, and CA3 regions on the probe was then estimated through manual inspection of LFP features and spike distribution as well as supplementary spectral analysis. Each recording day, the detailed recording configuration of channels was optimized (1) based on the location of hippocampal recording channels as estimated during the initial test recording, (2) by manual selection of channels exhibiting higher spike activity and (3) by inclusion of channels with the same x-coordinate and evenly spaced y-coordinates (40 μm apart) to facilitate subsequent current source density (CSD) analysis. The recording configuration was adjusted daily to achieve optimal yields.

### Spike sorting

Data recorded from the Neuropixels 1.0 probe was first preprocessed using the CatGT tool (https://billkarsh.github.io/SpikeGLX/help/dmx_vs_gbl/dmx_vs_gbl) and then spike sorted with Kilosort 2.5 (https://github.com/MouseLand/Kilosort). Manual curation of sorted clusters was performed in Phy (https://github.com/cortex-lab/phy) to remove non-neuronal clusters and assess cluster quality using two metrics: contamination index and amplitude cutoff. The contamination index, a default output by Kilosort 2.5, reflected the degree of spike contamination in a cluster. It was calculated as the ratio of spike rates occurring within refractory periods to those outside and ranged from 0 to infinity. The amplitude cutoff estimated the proportion of missed spikes in a cluster, assuming spike amplitude follows a Gaussian distribution and ranged from 0 to 0.5 (https://github.com/AllenInstitute/ecephys_spike_sorting) ^71^. Clusters with a contamination index below 0.8 and an amplitude cutoff below 0.5 were selected for single-unit and spike decoding analysis based on the experimenters’ manual inspection experience. Putative excitatory and inhibitory cells were distinguished by spike width (principal cells > 0.5 ms; interneurons < 0.5 ms), and the number of selected cells for each neuron type is detailed in Table S2.

### Instantaneous velocity

The rat’s x and y position (sampling rate 30 Hz) on the radial maze was respectively smoothed using a 2^nd^ order low pass digital Butterworth filter with a cutoff frequency of 1 Hz. For each sampling time interval (1/30 s), the rat’s instantaneous velocity was calculated as distance traveled using the smoothed position data on the radial maze divided by the interval length.

### Current source density (CSD) analysis

CSD analysis was performed using hippocampus channels with the same x-coordinate and evenly spaced y-coordinates (40 μm difference between adjacent y-coordinates) on the Neuropixels 1.0 probe. LFP data from these channels were downsampled to 625 Hz for the subsequent analysis and filtered between 5 and 100 Hz for CSD computation. Standard CSD analysis ^72^ was applied to the filtered data using Hamming’s three-point formula for noise reduction.

### Dorsal hippocampus layer estimation

Seven dorsal hippocampus layers were estimated by comparing the probe locations reconstructed from serial histology sections with LFP signal features from recording sites along the probe. The CA1 stratum pyramidale (sp), stratum radiatum (sr), and stratum lacunosum moleculare (slm) were identified by the channel with the highest LFP ripple power (150-250 Hz), the largest sink value in the CSD during ripple oscillations detected in CA1 ^73^, and the largest negative LFP amplitude during CA1 ripples, respectively. Dorsal hippocampus channels deeper than the CA1 slm were estimated to be in the DG-CA3 subregion. The outer molecular layer (oml), middle molecular layer (mml), and granule cell layer (gcl) in the DG were sequentially estimated by three major sink locations on the probe, based on CSD analysis during dentate spikes in the DG-CA3 area ^74, 75^. The CSD in the DG-CA3 area often appeared symmetric, as the Neuropixels 1.0 probe passed through the upper and lower blades of the DG, either fully or partially, allowing the CA3/hilus to be estimated as the channels at the center of the symmetric CSD pattern.

### Neural spike train decoding

Qualified clusters from dorsal hippocampus recordings (see Table S2 for cluster numbers and hippocampal subregions in each of the 16 sessions) were used to reconstruct an animal’s position using a Bayesian probability algorithm with a point process filter ^42–44, 76^. Briefly, at time stamp *t*, given a spike train *spike*_*t*_ from *N* sorted units, the posterior probability of the animal’s position *x*_*t*_ was calculated as:

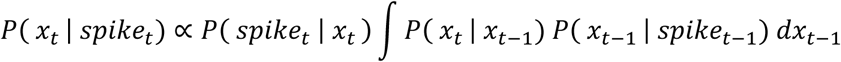

where

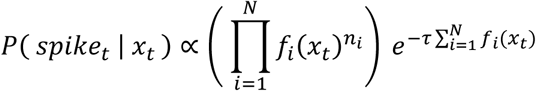

*f*_*i*_(*x*_*t*_) and *n*_*i*_ represent the firing rate and spike count of the *i*^*t*ℎ^ unit at position *x*_*t*_ within the decoding window *τ*, respectively. These calculations assumed (1) the activity of all *N* units to be independent, following Poisson firing statistics and (2) the animal’s position *x*_*t*_ to depend on its previous position *x*_*t*−1_. For simplicity, position data were binned into 2 cm by 2 cm bins. The variable *f*_*i*_(*x*_*t*_) was estimated from each unit’s tuning curve on the maze, which was calculated by dividing the number of spikes associated with each position bin by the time the animal spent in that bin when the animal’s moving speed exceeded 10 cm/s and by smoothing the raw rates by a 2D rotationally symmetric Gaussian low-pass filter of size 60 bins (120 cm) with standard deviation of 2 bins (4 cm). The position transition matrix *P*(*x*_*t*_ | *x*_*t*−1_) was defined as the probability of the rat occupying position *x*_*t*_ given the previous position *x*_*t*−1_ and was estimated as a constant numerical matrix based on position data from the entire maze session, also under the same velocity restriction (speed >10 cm/s). Decoder performance was measured by calculating the decoding error with 250 ms decoding windows and is reported for each of the 16 sessions in Table S2. Position reconstruction during theta states was performed in overlapping 20 ms windows with 5 ms advancement. A heatmap that displayed the probabilities of decoded locations over the maze was generated, and for further analysis, the summed probabilities for each of the eight arms and for the center platform were calculated. The region of the maze with the highest decoded probability, either one of the eight arms or the center platform, was identified as the maximally represented zone for each decoding window.

### Definition of behavioral epochs

The maze was divided into eight individual arms and the center platform. The center platform was defined as the area where the distance to the center of the maze was no greater than 30 cm. The arms were defined as areas on the individual arms where the distance to the center was greater than 30 cm. Reward zones were defined as the ends of the arms, located more than 70 cm from the center. Rat behavior during each trial was divided into four zones based on the animal’s position, and in some cases, restricted by velocity: (1) Occupying the center platform at any velocity, (2) Performing an outbound run in the arm (i.e., from the center platform toward the arm end) with a velocity ≥10 cm/s, (3) Occupying the reward zone at any velocity (which typically began with the consumption of chocolate milk in correct visits, was followed by varied behaviors that included prolonged pauses, and concluded with reorientation of body direction towards the inbound run), and (4) Performing an inbound run in the arm with a velocity ≥10 cm/s (i.e., from the arm end to the center platform).

### Spectral analysis

The LFP spectrum in hippocampus layers during the spatial working memory task as well as during pre- and post-maze sleep box sessions was calculated in 2-s windows using a multi-taper spectral analysis toolbox (http://chronux.org/) ^77^. The theta to delta power ratio was computed based on filtered LFP power in the delta (1-4 Hz) and theta (6-12 Hz) bands and averaged in 2-s time windows.

### Sleep periods

Sleep periods were defined as extended periods of immobility, when 2-s windows indicated immobility (i.e., at least 95% of timestamps within each window <1 cm/s) and consecutive immobility windows lasted over 50 seconds in pre- and post-task sleep box sessions. Within these identified sleep epochs, rapid eye movement (REM) sleep was defined as periods with an average theta/delta LFP power ratio greater than 1 in 2-s windows, lasting more than 10 seconds. Remaining sleep periods were classified as slow-wave sleep. Identified sleep epochs for individual sessions are detailed in Table S1.

### Definition of theta periods

The LFP signal from DG oml was selected as the reference channel for all theta-related analyses since this layer consistently demonstrated the highest theta oscillation power in the hippocampus (Fig. 1c and e). Specifically, the LFP was band-pass filtered between 6-12 Hz, and the theta phase was defined as the phase of the Hilbert transform of the filtered signal, with 0°/360° representing the trough of the filtered oscillation in DG oml. Individual theta cycles were separated by theta troughs, and additional criteria were applied to include qualified theta cycles: (1) cycle duration between 60 ms and 200 ms, (2) peak theta power higher than the mean theta power of the entire recording session, (3) theta phase monotonically increasing during the cycle, with a phase range greater than 330° and (4) theta cycles not partially or fully overlapping with CA1 SWRs.

### Theta modulation of single units

For each qualified cluster, every spike was assigned to the nearest theta phase measured from the LFP in DG oml. Firing rates in theta phase bins (bin size: 10°) were calculated and smoothed using a Gaussian filter (standard deviation = 1.2) separately for four velocity ranges (0-1 cm/s, 1-5 cm/s, 5-10 cm/s, and >10 cm/s). Units with fewer than 30 spikes during a velocity condition were excluded from further analyses to ensure sufficient sampling. Theta-phase locking of hippocampal single units was identified using the circular Rayleigh’s test for uniform distribution of spikes in theta oscillations, with a significance level of 0.05, employing the CircStat Matlab toolbox ^78^. The theta modulation index for each unit was calculated from theta phase bins as (peak_rate-min_rate)/peak_rate, yielding values between [0, 1]. A higher theta modulation index indicates greater theta-phase locking firing for a cell. The peak phase of each CA1 interneuron, as well as of each DG/CA3 principal cell and interneurons, was defined as the phase bin with the maximum rate. To account for the bimodal phase distribution of CA1 principal neurons (CA1 PN; Fig 2j), a major and a minor peak were identified for CA1 PN. The major peak phase of each CA1 PN was defined as the theta phase bin with the maximum rate within [70°, 280°], while the minor peak phase was defined as the theta phase bin with the maximum rate in the remaining phases of the theta cycle ^16^.

### Theta sequence analyses

#### Maximally represented region in theta sequences

Analysis of information content of theta sequences focused on identifying the maximally decoded region of the maze within each theta cycle. As detailed above (see Neural spike train decoding) decoded probability maps were initially generated for each 20 ms decoding window. Decoding windows containing no spikes were excluded, and for windows containing at least one spike from qualified clusters, the theta phase (from the LFP in DG oml) closest to the center of the window was assigned. To ensure adequate sampling within each theta cycle, only theta cycles containing at least 14 decoding windows were included (included proportion of theta cycles: 97.9% ± 3.2%, mean ± sd, *n* = 16 sessions). The primary represented region (either center platform or one of 8 arms) within each theta cycle was defined as the zone that was most frequently represented across all decoding windows within the cycle. Theta cycles with dispersed maze representations— defined as a combined proportion of decoding windows representing the center platform and the maximally represented arm <0.5—were excluded (included proportion of theta cycles: 97.5% ± 0.62%, mean ± sd, *n* = 16 sessions). Theta sequences were defined as local/remote if the maximally represented region (either center platform or one of 8 arms) corresponded/did not correspond to the rat’s currently occupied region.

#### Average theta sequences

The average theta sequence analysis was used to quantify the average represented pattern of a group of theta sequences. First, a standard approach for average theta sequence analysis was fused to quantify the movement theta sequences during inbound/outbound journeys (i.e., Fig. S8). Here the average spatial representations with reference to the position of the rat (in 2 cm bins, along the axis from the center to the end of the arm) was calculated for each phase bin (10°) in each qualified theta cycle. Thus, each coordinate (x, y) shows the average decoding probability of relative position y at phase bin x. This approach was suitable for movement theta sequences, when representations were tied to the animal’s current position and movement direction, but needed modification for the analysis of immobility theta sequences when representations included remote locations. For the modified approach, average theta sequences were quantified as 2D theta phase-position distribution density heatmaps (i.e., Fig. 3b, d, and f, Fig. S9i), where each coordinate (x, y) reflected the probability density of position y being represented at phase bin x, with normalization applied vertically at each phase bin. To generate such a heatmap, pairs of consecutive theta cycles that both represented the center platform, both represented the same maze arm, or represented the center platform and one maze arm were included (included theta cycles: 64.7% ± 3.1%, mean ± sd, *n* = 16 sessions) to ensure analysis of at least one complete theta sequence. For each phase bin (10°) within each included theta cycle pair, the maximally represented position relative to the rat’s current position (in 2 cm bins, along the axis from the center to the end of the arm) was identified. Thus, a 2D theta phase-position map can be generated to summarize above data, where each coordinate (x, y) reflected the count of relative position y being represented at phase bin x. Further normalization by the sum across position bins at each phase bin gave the final 2D theta phase-position distribution density heatmap. For each session, such 2D decoding probability heatmaps were calculated separately for the four behavioral zones (reward, center platform, inbound, and outbound movement) and four velocity ranges (0–1 cm/s, 1–5 cm/s, 5–10 cm/s, and >10 cm/s). To ensure sufficient sampling and robust analysis, conditions and sessions with fewer than 50 theta cycle pairs were excluded from subsequent phase parsing analyses for theta sequences (i.e., onset and core phases).

#### Onset and core phases of theta sequences

The onset and core phases of theta sequences were identified as the theta phase bins in the 2D theta phase-distance distribution density heatmap of each condition and session that showed the lowest and highest maximal distribution density, respectively, within the [0°, 360°] phase range. Individual theta sequences in each condition were thus characterized by the decoding probabilistic representation starting from the onset phase and ending up to 360° later.

#### Detailed quantification of theta sequence properties

Individual theta sequences were categorized based on correspondence to velocity, the rat’s location in maze zones and local/remote coding. For detailed quantification, only theta sequences with spike decoding in at least 20 phase bins (1 phase bin = 10°) were included in the analysis (included proportion of theta sequences: 83.0% ± 12.6%, mean ± sd, *n* = 16 sessions). For each theta sequence representation, a weighted best-fit line was calculated using a least-squares solution to approximate the spatial path represented in the sequence. Posterior probabilities within 5 cm of this line were averaged per phase bin, and these averages were combined across all phase bins to compute a sequence score ^16, 79^. A higher sequence score indicated a greater overlap between the best-fit line and the decoding probabilistic representation. The statistical significance of the sequence score was evaluated using a Monte Carlo approach with 100 shuffled posterior probability iterations (circularly rotating the posterior probability along the position axis for each 10° theta phase bin). Theta sequences with sequence score *P* values ≤ 0.05 were considered significant (Fig. S9b). The theta sequence length was calculated as the total distance along the maximally represented arm and center platform as determined by the best-fit line (Fig. S9c). The slope of the best-fit line defined the directionality of the spatial trajectory: positive slopes indicated movement from the center platform to the arm end, whereas negative slopes indicated movement from the arm end toward the center platform (Fig. S9d).

#### Next arm prediction analysis

Brain states were defined by the presence or absence of defined oscillation activities, which included periods of prominent theta oscillations (as defined in Theta cycle analysis), SWR activity (as defined in Detection of CA1 SWRs, see below), or periods that lacked prominent theta or SWRs (Non-Theta-Non-SWR periods). Non-Theta-Non-SWR periods were defined as periods of velocity <10 cm/s excluding CA1 SWR events, during which the z-scored theta power in DG oml was below -1. Next, behavioral epochs were defined as time segments from entry into either one reward zone or the center platform to the exit from that zone (Fig. S10a). The maximally represented region (one of eight arms or the center platform) was identified for each decoding window, and the summed representing probability during each epoch was calculated as the proportions of represented regions over the decoding windows of each epoch. For this calculation, the summed representing probability was calculated separately for SWRs, theta periods and Non-Theta-Non-SWR periods, and theta and Non-Theta-Non-SWR period were further divided by velocity ranges. The representing probability of the next arm the rat moved to was then identified for each of the brain state/velocity conditions. To assess whether the next arm was represented above chance (Fig. S10b), shuffled arm candidates were defined as the 6 arms excluding both the next arm and the current arm (during reward conditions) or the next arm and the arm the animal had just left (during center platform conditions). In each iteration, a randomly selected shuffled arm was independently chosen from this pool for each behavioral epoch. Across all behavioral epochs from 16 sessions, the preference for the next arm above chance was calculated as the difference between the proportion of epochs where the next arm had a higher decoded probability than a shuffled arm and the proportion of epochs where it had a lower decoded probability. This shuffle process was repeated 500 times, and the mean of the 500 preference values was tested for being greater than zero using a one-sided *t*-test. A *P* value < 0.05 indicated that the next arm was represented above chance. In the reward condition, epochs were analyzed separately for the forced phase (1st to 3rd rewards) and the choice phase (4th to 7th rewards). Similarly, for the center platform condition, epochs following the 1st to 3rd rewards (forced phase) and the 4th to 7th rewards (choice phase) were analyzed separately. The corresponding analysis was performed by replacing the next arm with the arm immediately preceding the rat’s current arm (during reward conditions) or the arm the animal had just left (during center platform conditions; Fig. S11).

#### Correlation of next arm representation between reward and center pairs

The decoded probability of the next arm during periods of velocity <10 cm/s was analyzed for reward-center epoch pairs with at least 10 decoding frames each. Pearson correlation was used to assess the relationship between next arm representations in reward and subsequent center epochs. A positive correlation coefficient with a *P*-value < 0.05 indicated a significant positive correlation in next arm representation between the reward epoch and the following center epoch. Epochs during forced and choice phases were analyzed separately.

### Detection of CA1 SWRs

Qualified CA1 SWRs were defined as the co-occurrence of ripples in the CA1 pyramidal layer and sharp waves in the CA1 stratum radiatum ^67^. Ripples were detected as periods when the power of the filtered LFP in the ripple band (150-250 Hz, averaged across three adjacent center channels in the CA1 pyramidal layer) was higher than 3 standard deviations above the mean for at least 35 ms, using only periods of velocity <10 cm/s. The start and end of each ripple were defined as the points when the average ripple power crossed the mean ^5^. Sharp waves were identified as troughs (minimum of 2 standard deviations below the mean) in the filtered (bandpass filter 5-40 Hz) and smoothed (Gaussian, sigma 12.5 ms) LFP signal from the center channel in the CA1 stratum radiatum. Only ripple events that co-occurred with sharp waves were designated as CA1 SWRs for subsequent analysis. Average LFP traces of CA1 SWRs were aligned by the peak of the filtered LFP amplitude (ripple band: 150-250 Hz) in the pyramidal cell layer. For Fig. S12c and d, each awake SWR on the maze was matched to a randomly selected event (’matched Non-SWR period’) that fulfilled the following criteria. The matched period was outside of SWRs while the animal’s velocity was within ± 1 cm/s and LFP theta power was within ± 0.1 (z-score) of the corresponding SWR. The SWR and the matched Non-SWR periods were of the same duration and were aligned by matching the theta phase (± 5°} at the peak of the Non-SWR period to the theta phase at the ripple peak.

### Information content of CA1 SWRs

Spikes during CA1 SWRs were decoded using 20 ms decoding windows with a 5 ms advancement. The maximally represented region for each CA1 SWR was determined by identifying the area on the maze with the highest representing probability across all decoding windows included in the SWR event. Local events refer to CA1 SWRs during which the maximally represented region coincided with the animal’s current arm or center location, while remote events indicate CA1 SWRs during which the maximally represented region did not align with the rat’s currently occupied zone.

