## Supplementary Information for "Mental exploration of future choices during immobility theta oscillations"

- 3 Supplementary Videos (associated .mp4 files)
- Supplementary Video Legends
- 12 Supplementary Figures
- 2 Supplementary Tables

**SUPPLEMENTARY VIDEO LEGENDS**

**Video S1. Theta sequences during movement and immobility.** The video illustrates theta sequences observed during the working memory task as the rat took an outbound journey on arm 3 and then paused in the reward zone. **Left Panel**, Decoded position based on hippocampal spike data (heat map). The rat's x-y position is represented by the blue circle for comparison. **Right Panel**, Concurrent z-scored LFP signals from hippocampal layers, with the DG oml layer highlighted in red. **Stage 1, Outbound journey, movement theta, local representation.** During movement-associated theta oscillations, the decoded position swept from behind to ahead of the rat, reflecting local theta sequences. **Stage 2, Reward, non-theta, local representation.** Upon reaching the reward zone, theta power decreased significantly, and the decoded position became stationary around the rat. **Stage 3, Reward, immobility theta, local representation.** While the rat remained immobile, prominent theta oscillations reappeared. Sequential representations of locations on arm 3 (immobility theta sequences) emerge, sweeping from the reward zone back towards the center platform.

**Video S2. Mental exploration during immobility theta in the reward zone.** The video demonstrates mental exploration during the working memory task when the rat paused at the end of arm 3 after retrieving the reward. Theta oscillations remained prominent throughout the example period. **Left Panel**, Decoded position based on hippocampal spike data (heat map). The rat's x-y position is represented by the blue circle for comparison. **Right Panel**, Concurrent z-scored LFP signals from hippocampal layers, with the DG oml layer highlighted in red. **Stage 1, Reward, immobility theta, current arm representation.** The rat's current location was represented. **Stage 2, Reward, immobility theta, flexible remote representations.** Multiple arm choices were mentally explored through theta sweeps during immobility. **Stage 3, Reward, immobility theta, consistent remote representation of next arm choice.** Spatial representation in theta sequences focused on a remote arm (arm 5), which later became the rat's next choice. **Stage 4, Reward, immobility theta, flickering local and remote representations between current location and next choice.** Flickering representations moving between arm 5 and the current arm were observed.

**Video S3. Mental exploration during immobility theta in the center platform.** The video illustrates flexible mental exploration when the rat is paused at the center platform and theta oscillations remained prominent. **Left Panel**, Decoded position based on hippocampal spike data (heat map). The rat's x-y position is represented by the blue circle for comparison. **Right Panel**, Concurrent z-scored LFP signals from hippocampal layers, with the DG oml layer highlighted in red. **Stage 1, Center, immobility theta, remote representations of multiple arms.** During this period, flexible representations of multiple arm choices were represented in theta sweeps, demonstrating the rat's mental exploration of potential navigational paths.

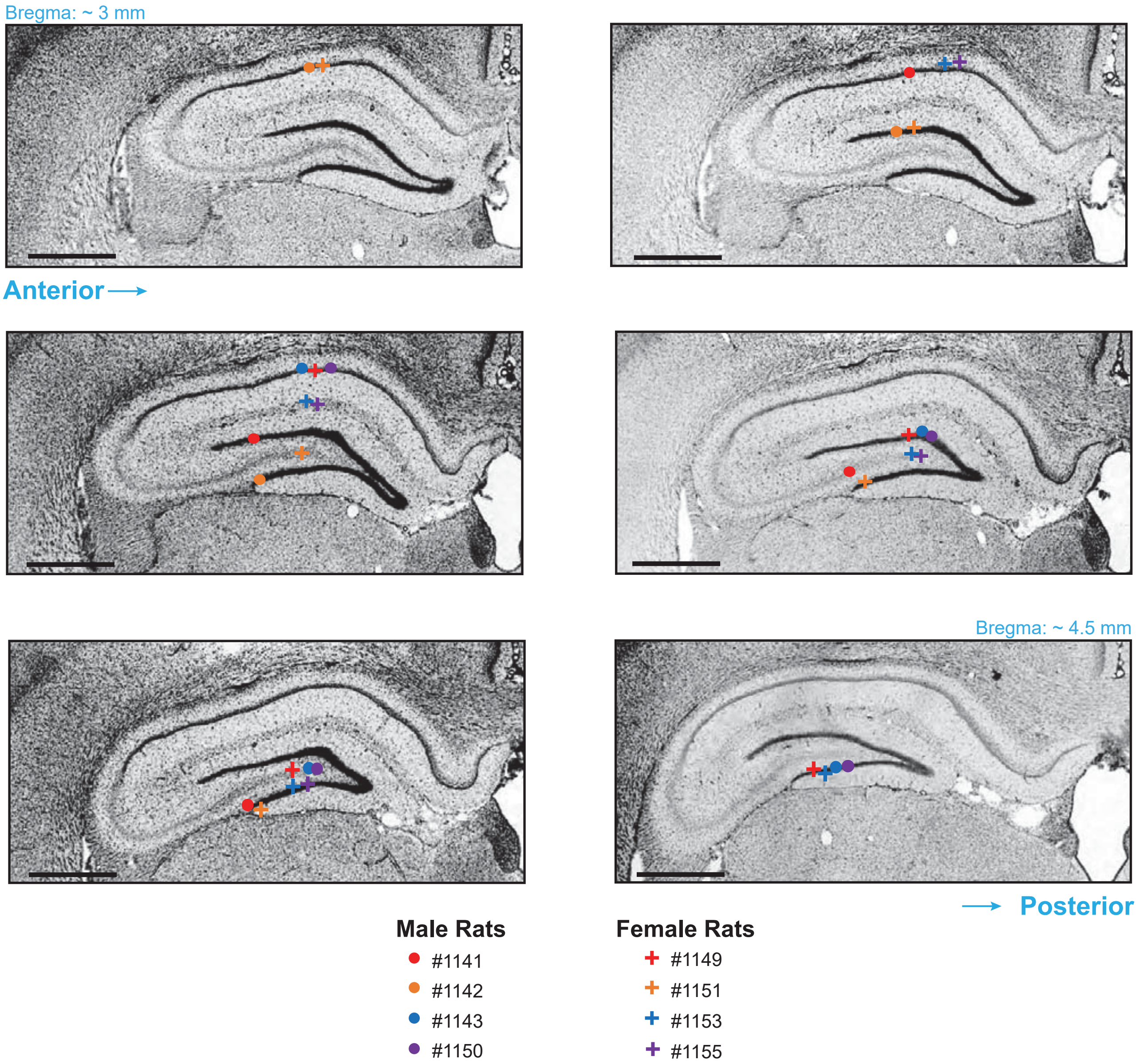


**Figure S1. Reconstruction of electrophysiological recording sites within the dorsal hippocampus.** Neuropixels 1.0 probes were dipped in DiI prior to implantation. Coronal serial sections were examined and the crossing of hippocampal cell layers identified with fluorescence microscopy. The location of probe tracks intersecting dorsal hippocampal CA1, DG, and CA3 layers for all experimental rats are superimposed on a series of sections from an example rat counterstained with DAPI (shown in black for visual reference). Coronal rat sections including the hippocampus are shown from anterior to posterior. Symbols of specific colors and shapes correspond to a probe location in one of 4 male rats (circles) and 4 female rats (crossed) included in the study. Each combination of color and symbol are specific to one rat. The identification number for each rat used in the study is shown here as well as in examples from individual rats in other figures, providing an easy reference for anatomical recording location in each instance. All CA3 recordings were located in the hilar/CA3c region. Recording locations were determined for analyses by comparing the anatomy, shown here, to the LFP and CSD profiles from multiple recording sites along the Neuropixels probe (see Methods). Scale bar, 1 mm.

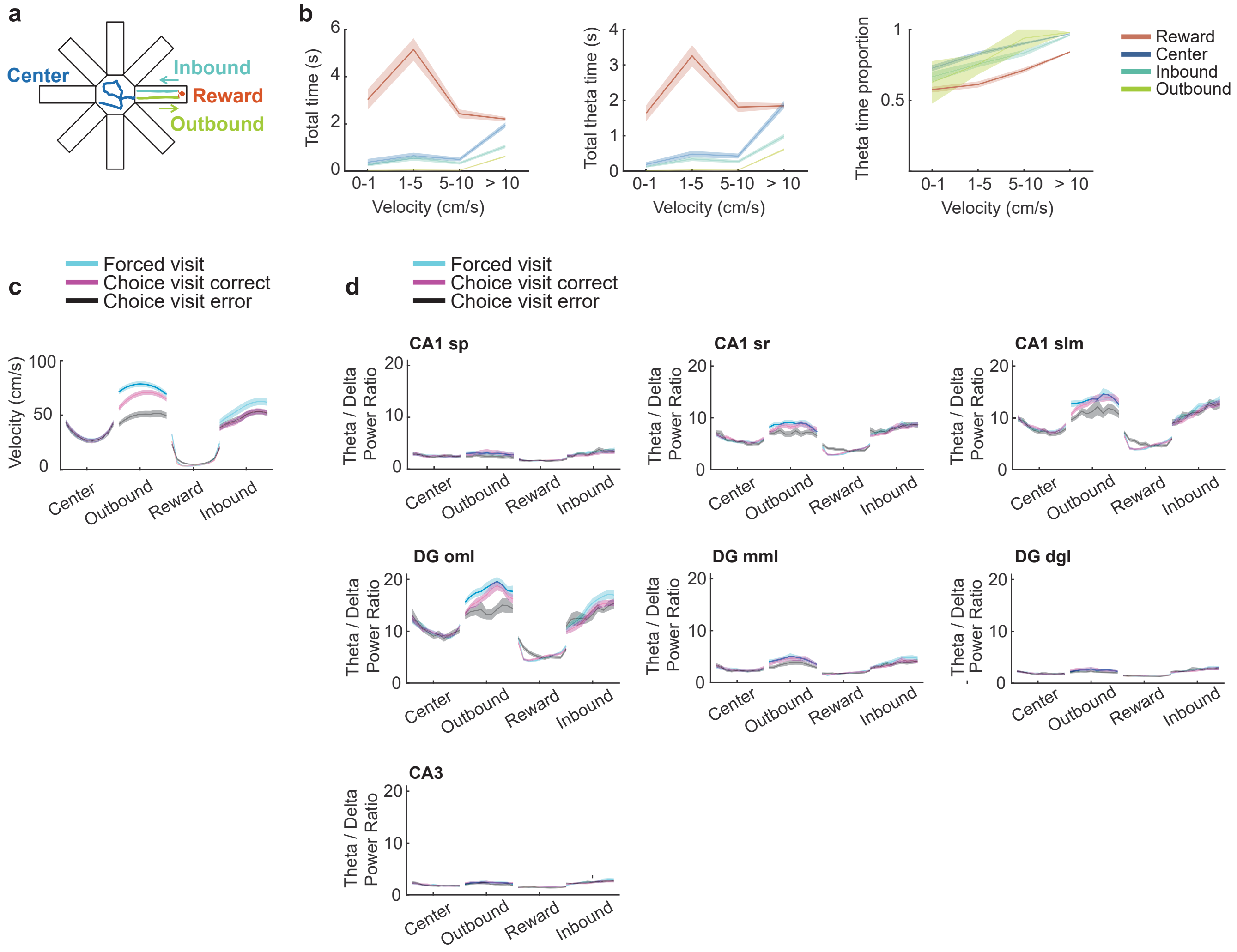


**Figure S2. Prominent theta oscillations were observed in the DG of the dorsal hippocampus when rats were immobile on the maze. a**, Schematic of how behavioral zones (reward, center, inbound and outbound) were defined by the animal's position on the maze and, for the middle of the maze arm, by the direction of travel (inbound and outbound in reference to the center). **b**, Total time (left), total theta state time (middle), and proportion of theta state time (right) as a function of the rats' velocity ranges, compared across the four defined behavior zones. Lines and shaded areas are mean ± s.e.m., *n* = 16 sessions. **c**, Mean velocity in each of the behavior zones on the maze, shown separately for arm visits in the forced and choice phase of the task. Arm visits during the choice phase were further subdivided into visits that were correct (arm had not yet been visited in the trial, Choice visit correct) or incorrect (arm had been visited earlier in the trial, Choice visit error). Average velocity was near zero when the rat was at the reward location. Lines and shaded areas are mean ± s.e.m., *n* = 16 sessions. **d**, Theta/delta power ratio for seven HPC layers as a function of behavior zones on the maze and compared across the different arm types (Forced visit, Choice visit correct, Choice visit incorrect). Theta oscillations were most pronounced in DG oml. Lines and shaded areas are mean ± s.e.m., *n* = 16 sessions. sp, stratum pyramidale; sr, stratum radiatum; slm, stratum lacunosum moleculare; DG, dentate gyrus; oml, outer molecular layer; mml, middle molecular layer; gcl, granule cell layer.

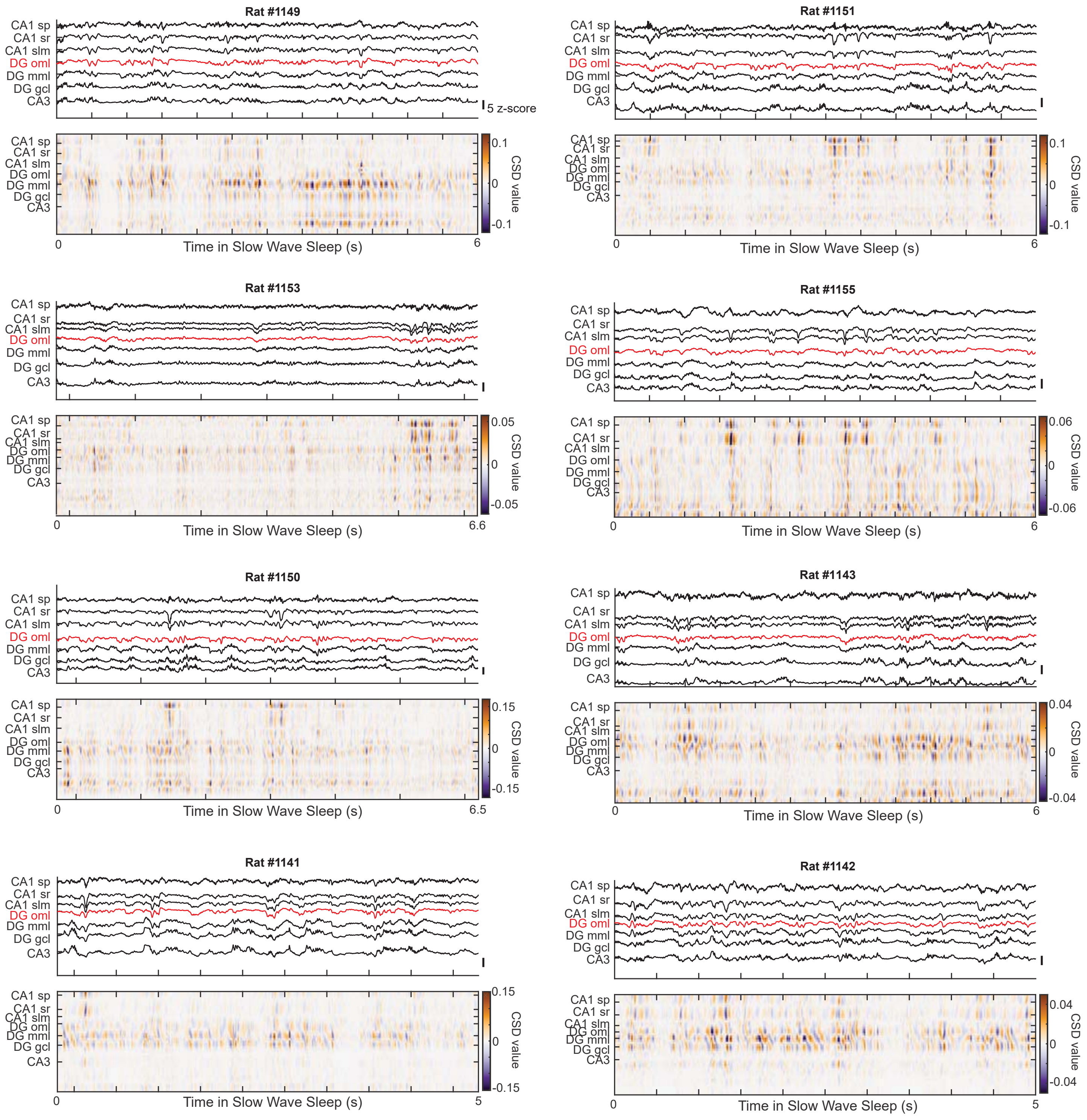


**Figure S3. Slow-wave sleep is characterized by CSD patterns that include SWRs.** Additional examples of LFP traces (z-scored, top of each panel) and the sinks (dark blue) and sources (dark orange) in CSD plots (bottom of each panel) during slow-wave sleep. Images are presented as described in Fig. 1f. Among the LFP traces, the trace from the HPC layer with the largest theta amplitude (DG oml) during behavior is labelled red for comparison. One example period is shown from each rat (*n* = 8 rats). LFP scale bars, 5 z-scores. sp, stratum pyramidale; sr, stratum radiatum; slm, stratum lacunosum moleculare; DG, dentate gyrus; oml, outer molecular layer; mml, middle molecular layer; gcl, granule cell layer.

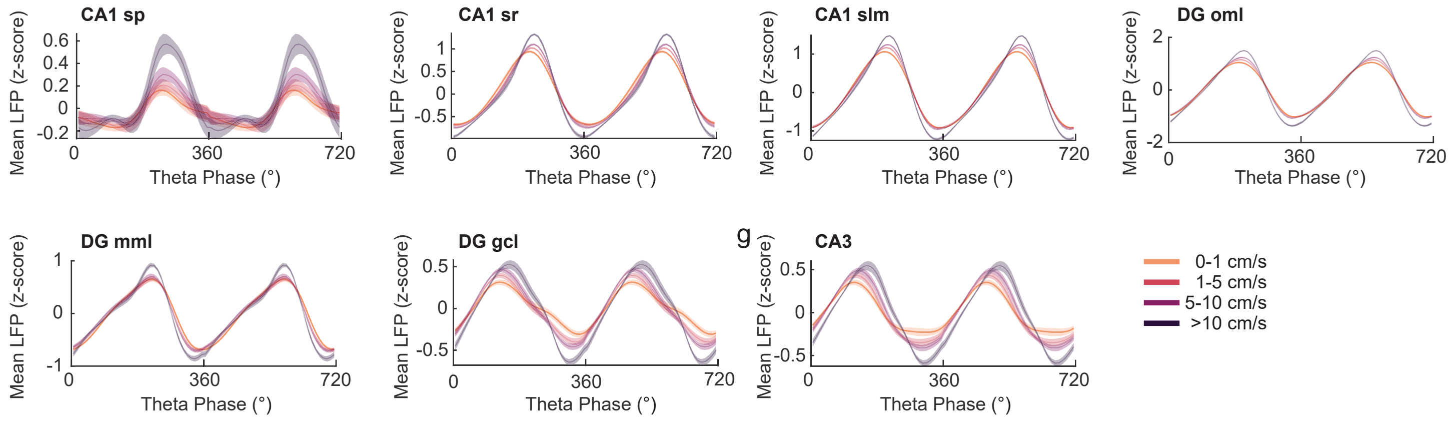


**Figure S4. Increasing asymmetry in theta waves with higher velocity.** Average LFP (z-scored) of theta cycles in seven HPC layers, compared across four velocity ranges. Note the increasing asymmetry in theta waveform with higher velocity. Lines and shaded areas are mean ± s.e.m., *n* = 16 sessions. sp, stratum pyramidale; sr, stratum radiatum; slm, stratum lacunosum moleculare; DG, dentate gyrus; oml, outer molecular layer; mml, middle molecular layer; gcl, granule cell layer.

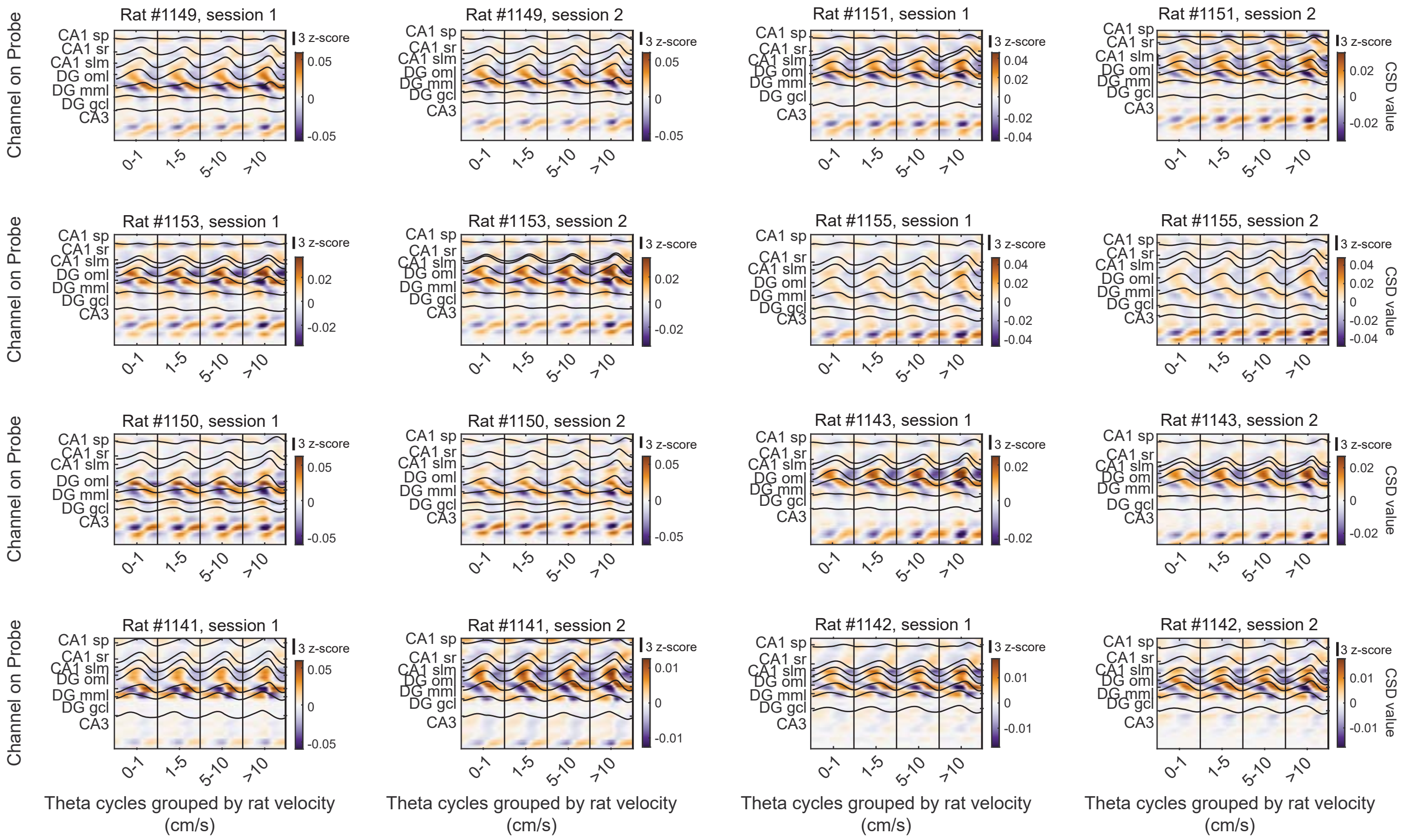


**Figure S5. Consistent theta-related sink and source patterns across HPC layers, irrespective of velocity during behavior.** Average CSD of theta cycles (maximum current sink, darkest blue; maximum current source, darkest orange) for four velocity ranges (0-1 cm/s, 1-5 cm/s, 5-10 cm/s, >10 cm/s) in 16 sessions (session and rat numbers on top of each panel). Average z-scored LFP traces from defined recording sites along the Neuropixels 1.0 probe (layers are specified on the y-axis) are overlaid (black traces). For individual animals and sessions, note the consistent CSD patterns of theta cycles across all velocity ranges.

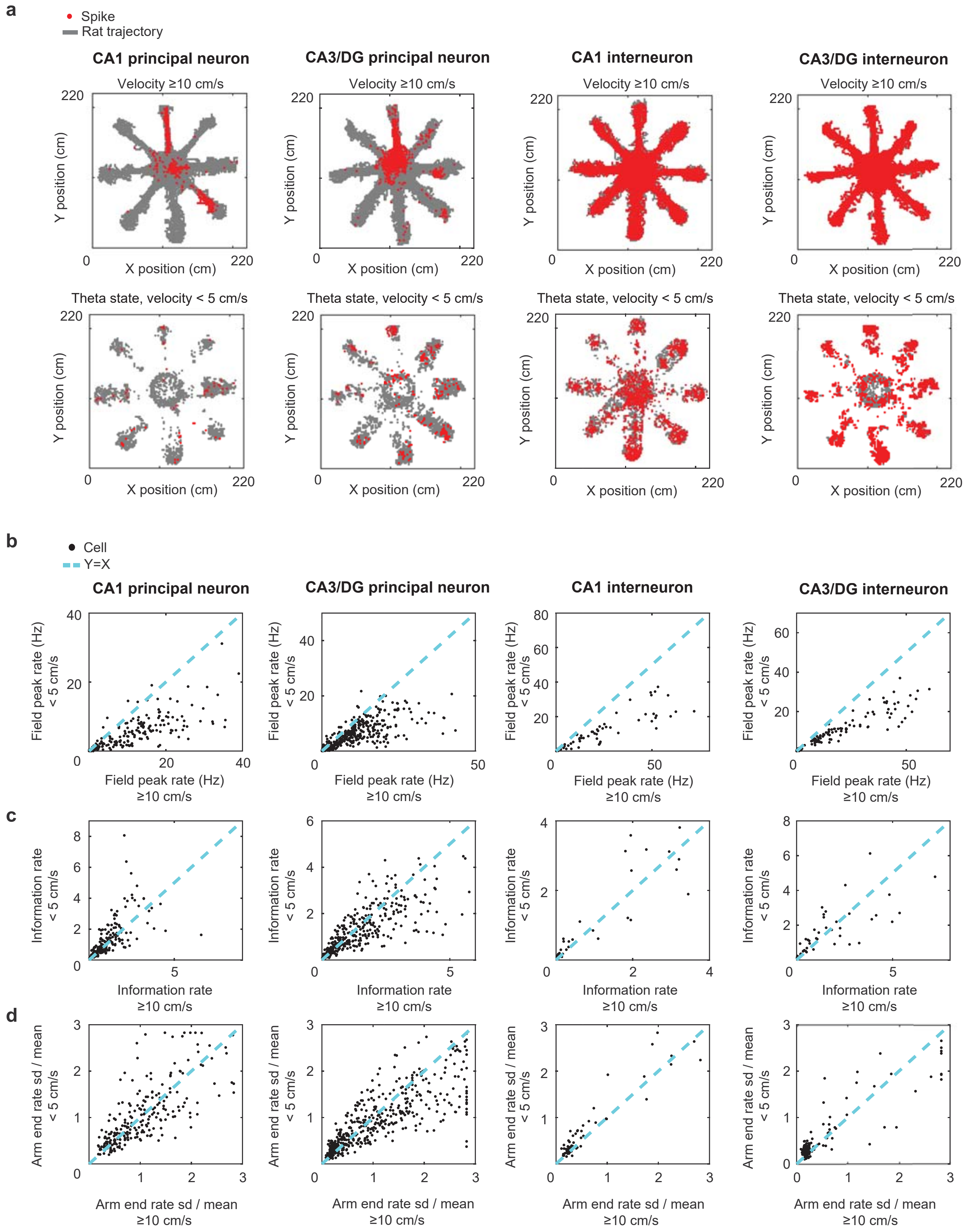


**Figure S6. Spatial firing features of hippocampal units compared between movement and immobility theta states in behavior. a**, Spatial distribution of spikes (red dots) overlaid on the rats' position traces (grey lines) for four example hippocampal units, categorized by cell type (CA1 principal neuron, CA3 principal neuron, CA1 interneuron and CA3 interneuron). Top row, spikes during movement velocity ≥10 cm/s. Bottom row, spikes from

**Figure S6 continued...**

the same neurons but during immobility theta states (velocity <5 cm/s). Out of place field firing is prominent with velocities under 5 cm/s. **b**, Comparison of peak firing rates for the four types of hippocampal cells during movement theta (x-axis) and immobility theta (y-axis). Each dot represents a single cell. Firing rates of the four cell types were higher during movement than immobility theta states, (from left to right, $t$ = 12.77, 14.74, 7.85, 10.11; $P = 4.4 \times 10^{-29}$, $3.6 \times 10^{-41}$, $3.9 \times 10^{-11}$, and $1.5 \times 10^{-18}$, one-sided $t$-tests; n = 239, 484, 62, 136 units, respectively). **c**, Information rate, displayed as in b. Information rate was higher during movement than immobility theta only for CA3 principal cells (from left to right, $t$ = -2.83, 4.47, -1.29, 0.012; $P$ = 1.0, $5 \times 10^{-6}$, 0.9, 0.5, one-sided $t$-tests). **d**, Reward-tuning specificity, displayed as in b. Tuning specificity was calculated as the standard deviation of firing rates at eight reward sites, normalized by the average firing rate at the reward sites and was higher during movement than immobility theta only for CA3 principal cells (from left to right, $t$ = -0.26, 5.37, -3.77, -1.40; $P$ = 0.6, $6 \times 10^{-8}$, 1, and 0.92, one-sided $t$-tests).

### Figure S7

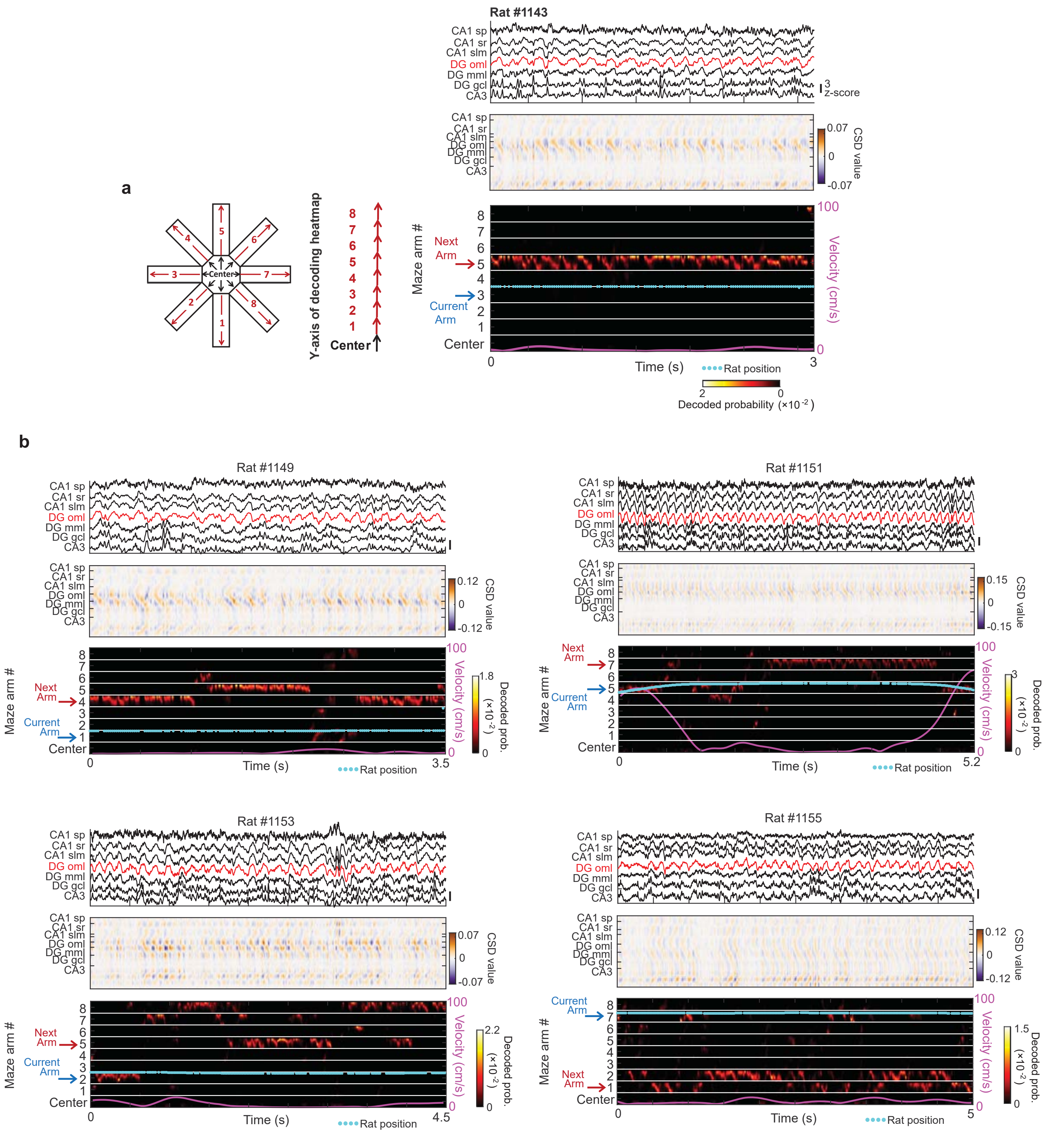

a
Rat #1143
CA1 sp
CA1 sr
CA1 slm
DG oml
DG mml
DG gcl
CA3
3
z-score
CSD value
0.07
0
-0.07
Y-axis of decoding heatmap
Center
Maze arm #
Next
Arm
Current
Arm
Velocity (cm/s)
100
0
Time (s)
Rat position
3
Decoded probability (×10⁻²)
b
Rat #1149
Rat #1151
Rat #1153
Rat #1155
0.12
-0.12
0.15
-0.15
1.8
3
2.2
1.5
Decoded prob. (×10⁻²)
3.5
5.2
4.5
5

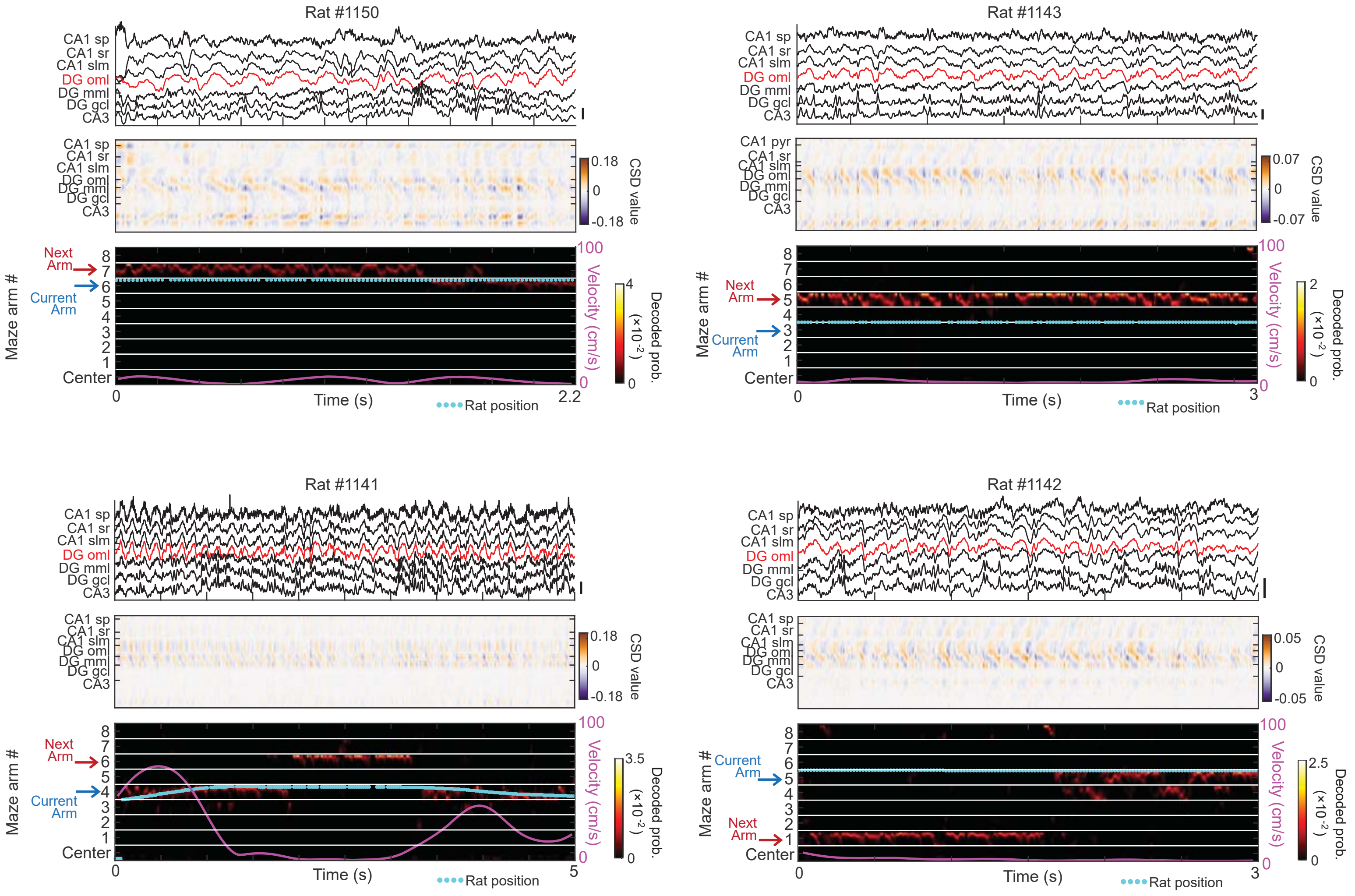


**Figure S7. Theta sequences during immobility in the reward zone preferentially represented the upcoming choice. a**, Example trial segment (3 s long), as shown in Fig. 3c, but here with a schematic (left) describing how the linearized heatmap that shows Bayesian decoded probability of hippocampal population activity (bottom panel) is organized with respect to the 8-arm maze. The y-axis of the heatmap represents linearized maze position with the center platform at the bottom of each bar and the 8 arms stacked in numerical order, not in the order of arm visits. Therefore, decoding probability on each maze arm is oriented with center platform positions close to the bottom white line and the reward zone close to the top white line of each arm segment. The current arm the rat explored during the segment is indicated by the blue arrow to the left of the y-axis, and the position of the rat on that arm at each time point is indicated by cyan dots. The next arm choice of the animal in the behavioral series is indicated by the red arrow to the left of the y-axis. The instantaneous velocity of the animal at each time point is overlaid (pink line) on the heatmap, using the scale to the right. The corresponding LFP traces from seven HPC layers (top) and CSD plots (middle) are shown for the same time segment. Among the LFP traces, the trace from the HPC layer with the largest theta amplitude (DG oml) during behavior is labelled red. **b**, Each panel shows one example trial segment during immobility in the reward zone for each rat in the study ($n$ = 8). All examples show representations by theta sequences that are biased toward the next arm the animal visited during the choice phase of the task. LFP scale bars, 3 z-scores.

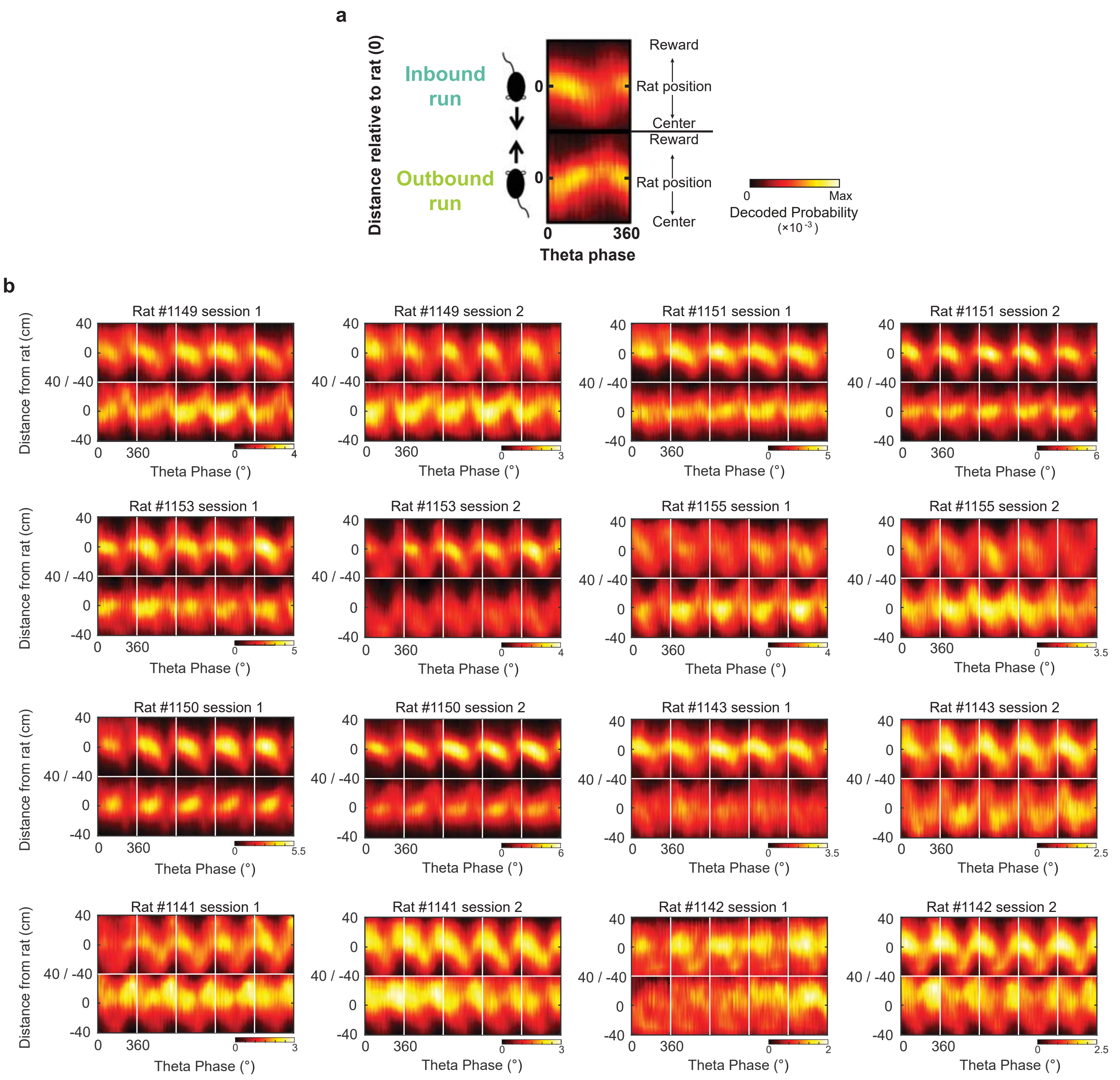


**Figure S8. Mean theta sequences during inbound and outbound journeys on maze arms. a**, Figure key. Heatmaps of represented locations on maze arms during inbound journeys (top) and outbound journeys (bottom). Each map is aligned to the current position of the rat (set at 0) and oriented such that the center of the maze is towards the bottom and the reward location towards the top. The x axis corresponds to a single theta cycle (0-360°, phase from DG oml). The pixel color is scaled to the maximum decoded probability (yellow).
**b**, Panels correspond to sessions (two per rat, *n* = 16 sessions). Top and bottom maps within panels are from inbound and outbound journeys, as illustrated in a. The columns of each top/bottom map display, from left to right, average theta sequences that were calculated from theta sequences with increasing velocity (starting at velocity ≥10 cm/s) within each session's inbound/outbound journeys. The total number of theta cycles for 16 sessions during inbound journeys are sequentially: 2066, 2589, 2811, 2000, 1721, 1774, 1889, 2041, 2594, 2402, 2301, 2246, 2642, 2937, 1882, 3077; during outbound journeys: 1259, 1722, 1744, 1254, 1174, 1323, 1233, 1579, 2427, 1700, 1428, 1347, 1714, 2123, 1399, 1951. Note that forward and backward hippocampal population representations of the space in movement theta sequences, as reported previously [16], are particularly evident during outbound runs in many sessions.

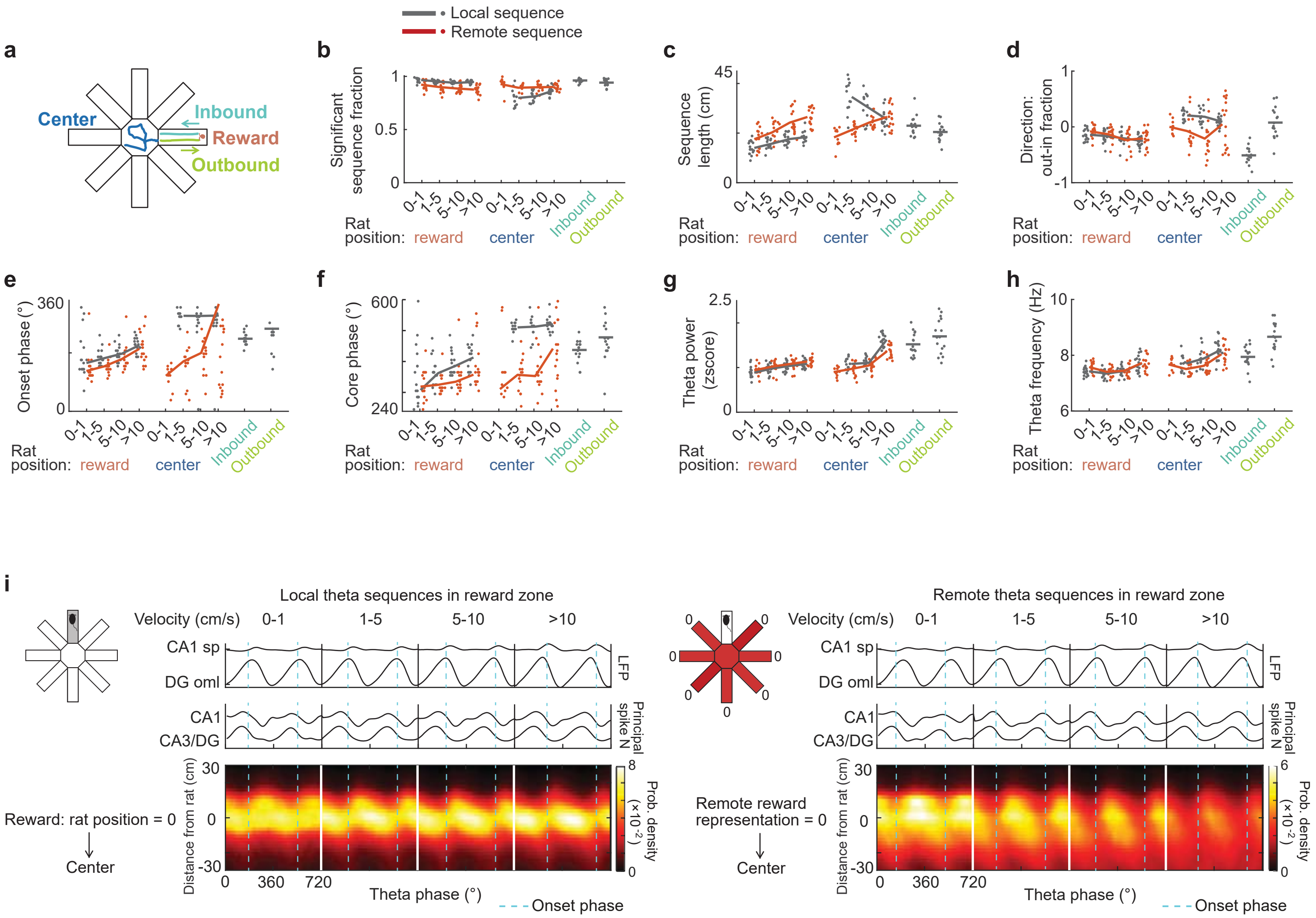


**Figure S9. Theta sequence properties differed depending on velocity range, behavior zone, and decoded content. a**, Segmentation of behavior zones (reward, center, inbound, and outbound) on the maze. Only movement periods (velocity ≥10 cm/s) were analyzed for inbound and outbound zones. **b**-**h**, Quantification and comparison of theta sequence properties across velocity ranges, behavior zones (as in a), and decoded content. For decoded content from hippocampal cell populations, we distinguished theta cycles when the maximally represented maze region by hippocampal activity matched the rat's current position on the maze (local theta sequence, gray) or a different region than the rat's current location (remote theta sequence, red; see methods for details). Sessions (*n* =16) with at least 50 qualified pairs of consecutive theta cycles in a condition (behavior zone/velocity range/decoded content) were analyzed and included in the figure (dots). For the center/0-1 cm $s^{-1}$/local condition, none of the sessions reached this criterion, and therefore, no data point is included. Dots are slightly jittered along the x-axis for improved visual clarity. Lines represent averages across sessions for each condition. b, The significant sequence fraction corresponds to the proportion of qualified theta sequences with continuous and smooth spatial trajectories, as opposed to erratic or jumping representations (see methods for details). The fractions of local compared to remote sequences differed in both the reward zone and at the center. Velocity only influenced the fraction in the reward zone, not at center platform (Reward: velocity, $F$ = 6.59, $P$ = 3.7×$10^{-4}$; local/remote coding, $F$ = 87.09, $P$ = 6.7×$10^{-16}$; interaction, $F$ = 1.09, $P$ = 0.36; Center: velocity, $F$ = 1.34, $P$ = 0.27; local/remote coding, $F$ = 30.73, $P$ = 1.9×$10^{-7}$; interaction, $F$ = 1.53, $P$ = 0.21, two-way ANOVAs, with velocity and local/remote coding as factors). **c**, The sequence length of theta sequences was measured as the spatial distance of represented virtual paths. Length of local and remote sequences in the reward zone or of

**Figure S9 continued...**

remote sequences in the center increased with higher animal velocity, whereas the opposite trend was observed for local sequences at the center (Reward: velocity, $F$ = 25.37, $P = 8.9\times10^{-13}$; local/remote coding, $F$ = 91.54, $P = 1.8\times10^{-16}$; interaction, $F$ = 2.97, $P$ = 0.034; Center: velocity, $F$ = 3.72, $P$ = 0.014; local/remote coding, $F$ = 43.82, $P = 1.2\times10^{-9}$; interaction, $F$ = 9.21, $P = 1.6\times10^{-5}$, two-way ANOVAs with velocity and local/remote coding as factors). **d**, Spatial trajectories of theta sequences can be either inbound (reward to center) or outbound (center to reward). The direction of a group of theta sequences was calculated as the difference in the proportion of outbound compared to inbound trajectories (see methods for details). Directional differences were observed between local and remote theta sequences at the center platform, but not in reward zone, and velocity had an effect on the directionality of theta sequences in both the reward zone and the center platform (Reward: velocity, $F$ = 6.69, $P = 3.3\times10^{-4}$; local/remote coding, $F$ = 1.77, $P$ = 0.19; interaction, $F$ = 0.74, $P$ = 0.53. Center: velocity, $F$ = 6.47, $P = 4.4\times10^{-4}$; local/remote coding, $F$ = 48.81, P = $2.0\times10^{-10}$; interaction, $F$ = 5.76, $P$ = 0.001, two-way ANOVAs, with velocity and local/remote coding as factors). **e**, Onset phases of local and remote theta sequences differed in the reward zone and at the center platform (Reward: velocity, $F$ = 15.80, $P = 1.0\times10^{-8}$; local/remote coding, $F$ = 11.6, $P = 8.9\times10^{-4}$; interaction, $F$ = 0.43, $P$ = 0.74; Center: velocity, $F$ = 6.67, $P$ = 0.15; local/remote coding, $F$ = 62.51, $P = 2.7\times10^{-14}$; interaction, $F$ = 3.66, $P$ = 0.16; parametric two-way ANOVAs for circular data, using the CircStat Matlab toolbox [78]). Onset phase of local and remote sequences positively correlated with velocity in the reward zone and at the center platform, except for local sequences at center platform (Reward: local coding, rho = 0.65, $P = 1.4\times10^{-6}$; remote coding, rho = 0.65, $P = 1.3\times10^{-6}$; Center: local coding, rho = 0.11, $P$ = 0.76; remote coding, rho = 0.43, $P$ = 0.0061, circular-linear correlations between onset phase and velocity levels, using the CircStat Matlab toolbox [78]). **f**, Core phases of local and remote theta sequences differed in the reward zone and at the center platform (Reward: velocity, $F$ = 32.33, $P = 1.4\times10^{-5}$; local/remote coding, $F$ = 24.27, $P = 5.4\times10^{-6}$; interaction, $F$ = 5.37, $P$ = 0.15; Center: velocity, $F$ = 6.93, $P$ = 0.14; local/remote coding, $F$ = 74.98, $P$ = 0.0; interaction, $F$ = 1.67, $P$ = 0.43; parametric two-way ANOVAs for circular data). Core phase of local and remote sequences positively correlated with velocity for reward and center epochs, except for local sequences at center platform (Reward: local coding, rho =0.54, $P = 7.5\times10^{-5}$; remote coding, rho = 0.34, $P$ = 0.025; Center: local coding, rho = 0.20, $P$ = 0.46; remote coding, rho = 0.34, $P$ = 0.04, circular-linear correlations between core phase and velocity levels). **g**, Local and remote sequences were associated with distinct theta power levels in the reward zone and at the center platform. Theta sequences of different velocity ranges were associated with distinct theta power levels in the reward zone and at the center platform (Reward: velocity, $F$ = 17.57, $P = 1.6\times10^{-9}$; local/remote coding, $F$ = 5.55, $P$ = 0.02; interaction, $F$ = 0.03, $P$ = 0.99; Center: velocity, $F$ = 76.35, $P = 2.0\times10^{-27}$; local/remote coding, $F$ = 7.08, $P$ = 0.0089; interaction, $F$ = 3.83, $P$ = 0.012, two-way ANOVAs, with velocity and local/remote coding as factors). **h**, At the center platform, theta frequency was higher during local compared to remote theta sequences, whereas in the reward zone, theta frequency during local and remote theta sequences did not differ. Theta sequences of different velocity ranges were associated with distinct theta frequency in the reward zone and at the center platform (Reward: velocity, $F$ = 16.21, $P = 6.6\times10^{-9}$; local/remote coding, $F$ = 2.72, $P$ = 0.10; interaction, $F$ = 0.81, $P$ = 0.49; Center: velocity, $F$ = 18.5, $P = 6.8\times10^{-10}$; local/remote coding, $F$ = 4.49, $P$ = 0.036; interaction, $F$ = 1.43, $P$ = 0.24, two-way ANOVAs, with velocity and local/remote coding as factors). **i**, Average theta sequence density heatmaps while rats were in the reward zone are shown separately for local coding (left) and remote coding (right). These plots, therefore, correspond to the bottom and top sections of the heatmaps in Fig. 3d respectively, but were normalized to their own phase bins. Theta cycles are sorted into four velocity ranges (the number of theta cycle pairs in each range, from left to right, local: 409.4 ± 45.8, 1445.3 ± 117.7, 986.5 ± 93.4, 1345.1 ± 172.3; remote: 1158.1 ± 163.9, 2512.8 ± 240.3, 1378.5 ± 116.1, 1112.4 ± 75.4, mean ± s.e.m., $n$ = 16 sessions). For local sequences, zero on the y-axis reflects the current position of the rat. For remote sequences, zero on the y-axis reflects the center position of the remote reward representation. Note that local and remote theta sequences at reward predominantly coded for inbound sweeps (i.e., from the end of the arm towards to the center).

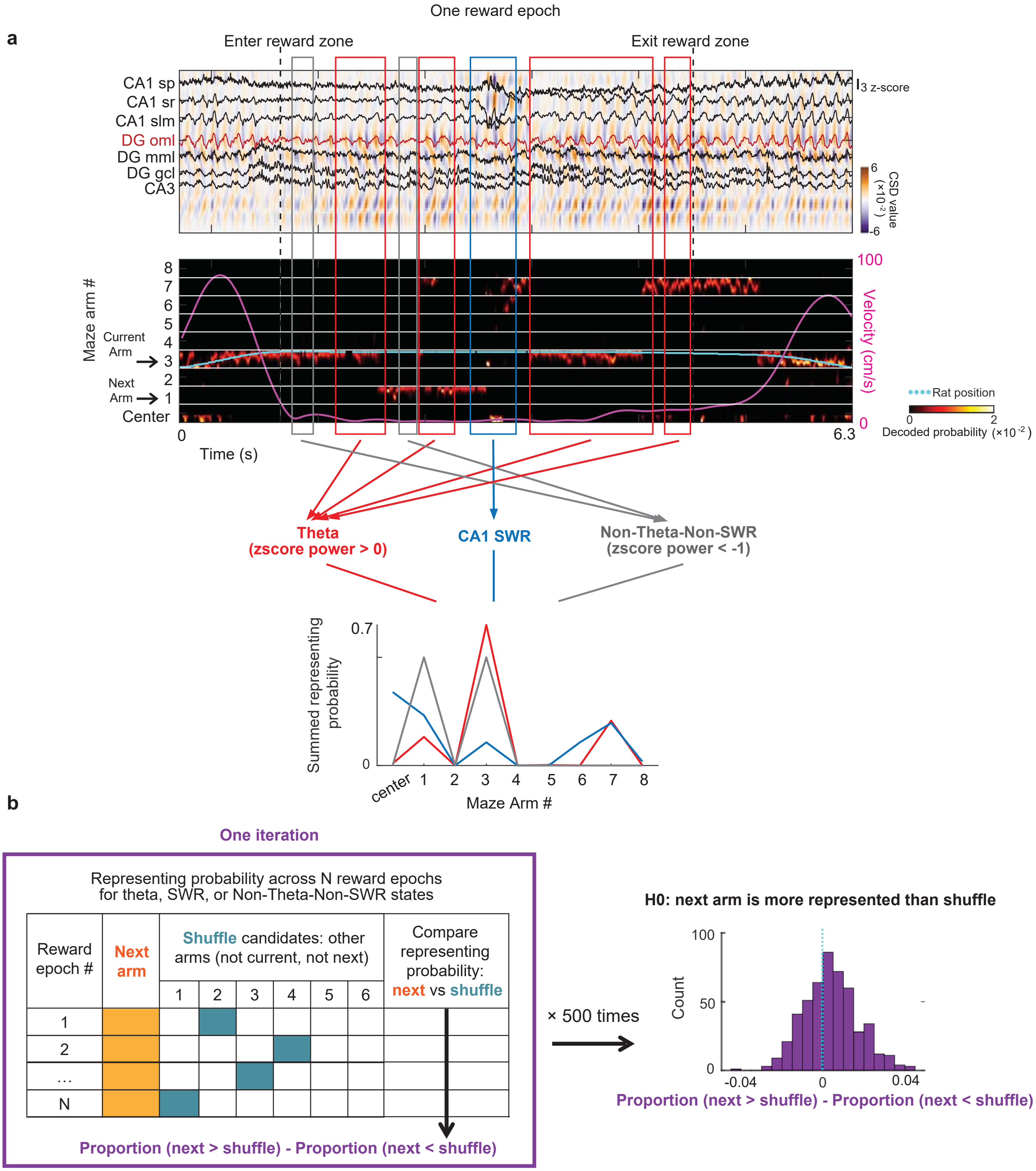


**Figure S10. Methods illustration for the next arm prediction analysis performed in Fig. 4a. a**, Throughout each epoch (from entry to exit from reward/center zone), brain states were classified as Theta (red), CA1 SWR (blue), and Non-Theta-Non-SWR (gray). Non-Theta-Non-SWR periods were defined as periods of velocity <10 cm/s excluding CA1 SWR events, during which the z-scored theta power in the DG oml was below -1. For each combination of state and velocity range, decoded probabilities of representing the eight arms and center platform were summed over each epoch. **b**, The decoded probability of the next arm the rat moved to was identified for

**Figure S10 continued...**

each of these conditions. To assess whether the next arm was represented above chance, shuffled arm candidates were defined as 6 arms excluding both the next arm and the current arm (during reward conditions) or the next arm and the arm the animal had just left (during center platform conditions). In each iteration, a randomly selected shuffled arm was independently chosen from this pool for each behavioral epoch. Across all behavioral epochs from 16 sessions, the preference for the next arm above chance was calculated as the difference between the proportion of epochs where the next arm had a higher decoded probability than a shuffled arm and the proportion of epochs where it had a lower decoded probability. This shuffle process was repeated 500 times, and the mean of the 500 preference values was tested for deviation from zero using a *t*-test. A mean greater than zero and a $P$ value $<0.05$ indicated that the next arm was represented above chance. In the reward condition, epochs were analyzed separately for the forced phase (1st to 3rd rewards) and the choice phase (4th to 7th rewards). Similarly, for the center platform condition, epochs following the 1st to 3rd rewards (forced phase) and the 4th to 7th rewards (choice phase) were analyzed separately. A corresponding analysis was performed by replacing the next arm with the arm immediately preceding the rat's current arm (during reward conditions) or the arm the animal had just left (during center platform conditions).

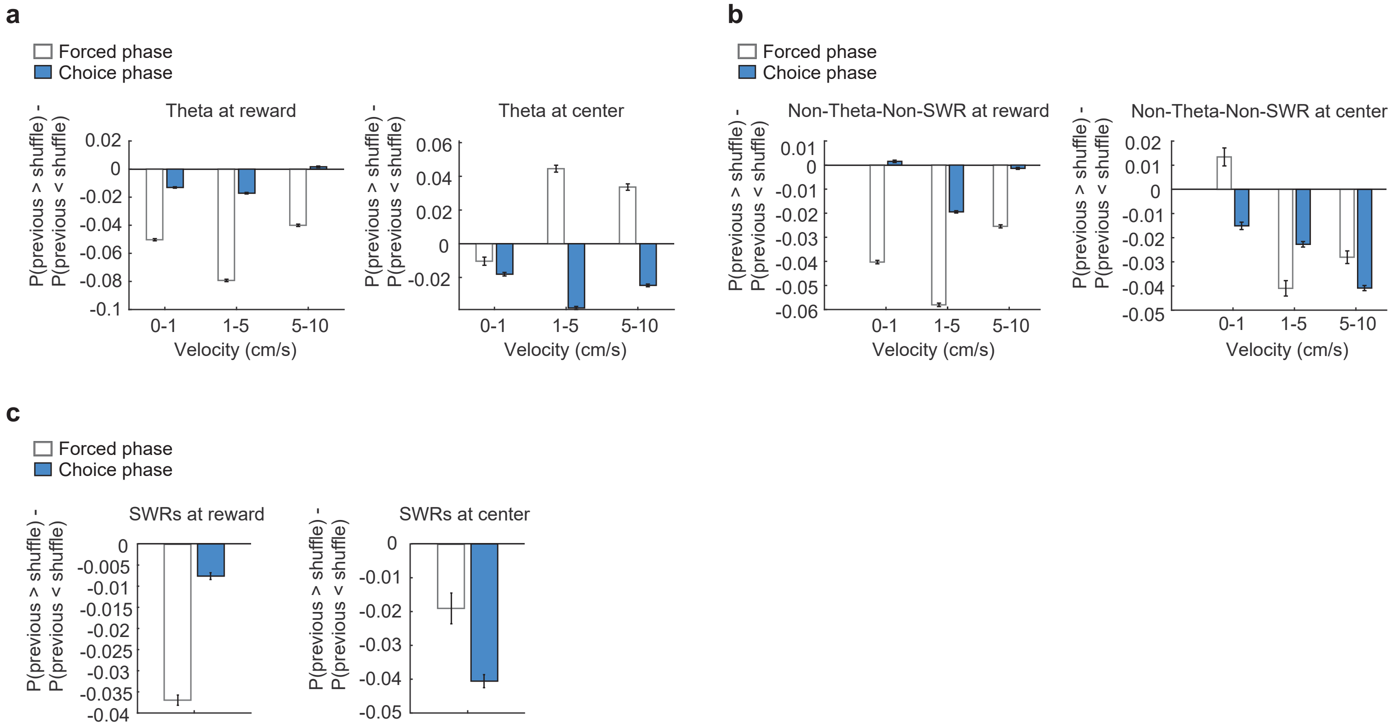


**Figure S11. Representations of the previous arm during theta and SWR states were less prominent than that of the next arm choice.** Analysis was performed as for the next arm choice in Fig. 4a, c, h, but by replacing the next arm choice with the previous arm choice. For reward zone analysis, this was the arm visit immediately preceding the rat's current arm visit, and for the center zone analysis, this was the arm the animal had just left. The coding preference for the previous arm was calculated as the difference between the proportion of epochs when the previous arm had a higher decoded probability than shuffled arms and the proportion of epochs when the previous arm had a lower decoded probability than shuffled arms ($n$ = 500 shuffles). Combinations of task phases (forced/choice), brain states (Theta, SWR, and Non-Theta-Non-SWR), and velocity ranges were analyzed and plotted separately. Velocity ranges were not distinguished for SWRs due to low occurrence rate. **a**, During theta states, coding for the previous arm was only higher than chance with the rat on the center platform during the forced phase (Reward: forced phase, $t$ = -64.09, -92.26, -48.11; $P$ = 1.0, 1.0, 1.0 for the three velocity ranges; choice phase, $t$ = -25.94, -30.64, 2.99; $P$ = 1.0, 1.0, 0.0014; Center: forced phase, $t$ = -4.28, 21.75, 17.7; $P$ = 1.0, $1.4\times10^{-74}$, $4.7\times10^{-55}$; choice phase, $t$ = -17.13, -42.43, -30.68; $P$ = 1.0, 1.0, 1.0, one-sided one sample $t$-tests). **b**, During Non-Theta-Non-SWR state, previous arm representations were not higher than chance, except in two combinations, reward zone/0-1 cm $s^{-1}$/choice phase and center/0-1 cm $s^{-1}$/forced phase (Reward: forced phase, $t$ = -56.31, -80.83, -38.65; $P$ = 1.0, 1.0, 1.0 for the three velocity ranges; choice phase, $t$ = 3.07, -40.49, -3.34; $P$ = 0.0011, 1.0, 1.0; Center: forced phase, $t$ = 3.61, -12.83, -10.75; $P$ = 0.00017, 1.0, 1.0; choice phase, $t$ = -9.89, -20.05, -38.41; $P$ = 1.0, 1.0, 1.0, one-sided one sample $t$-test). **c**, In SWRs, coding for the previous arm was not higher than coding for shuffled arms (Reward: forced phase, $t$ = -30.45; $P$ = 1.0; choice phase, $t$ = -9.48; $P$ = 1.0; Center, forced phase, $t$ = -4.19; $P$ = 1.0; choice phase, $t$ = -21.02; $P$ = 1.0, one-sided one sample $t$-tests). In contrast to next arm coding (Fig. 4a, h), previous arm coding was not prominent in theta and SWR states during the choice phase.

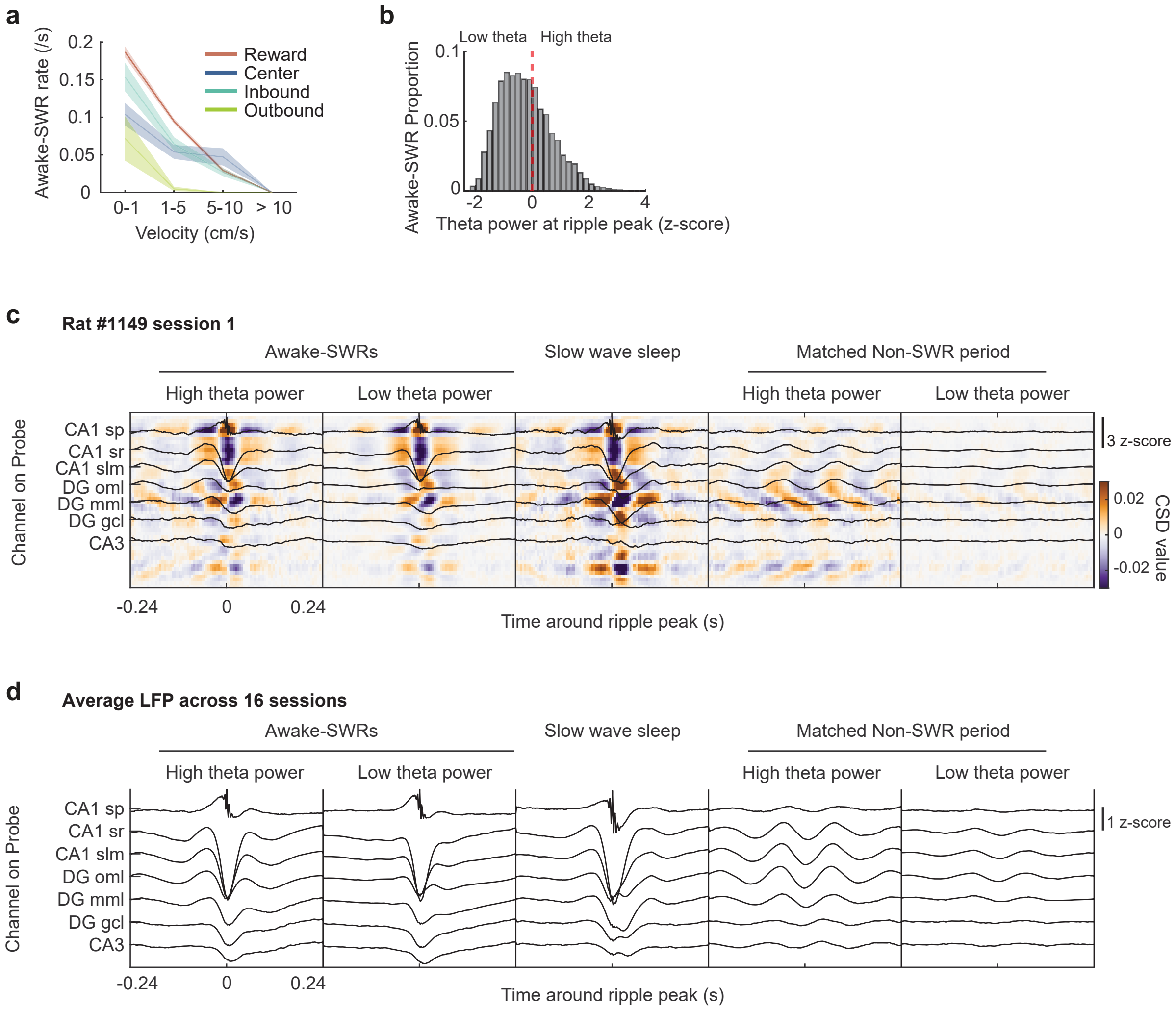


**Figure S12. Theta oscillations did not co-occur with SWRs during immobility. a**, Awake SWR event rate as a function of velocity ranges across behavioral zones. **b**, Distribution of z-scored LFP theta power aligned to the CA1 ripple peak time. Ripple events that coincided/did not coincide with high power in the theta band were defined by z-scored theta power larger/smaller than zero. Note that SWR-associated theta power can result from the sharp wave component rather than from sustained theta oscillations. **c**, Average CSD and overlaid LFP traces for SWRs on the maze, in sleep, and for matched Non-SWR periods on the maze are shown for an example session. Each SWR on the maze was matched to a period on the maze that did not overlap with a SWR ('matched Non-SWR period'), but when the rat's velocity and LFP theta power were approximately corresponding. In addition, matched Non-SWR periods (randomly selected from candidate periods) were of the same length as the corresponding SWRs and centered at the same theta phase as the ripple peak. SWRs were aligned by the ripple power peak and matched Non-SWR periods were aligned by their peak time. The CSD patterns corresponded between SWRs during sleep and behavior and between SWRs that were associated with high or low theta power. The CSD pattern of SWRs differed from matched Non-SWR periods, which resembled CSD profiles of theta states (see Fig. 2e). **d**, Average LFP traces (z-scored, *n* = 16 sessions) for the oscillation types described in c. The LFP profiles of awake SWRs that were either associated or not associated with high theta power were similar to that of SWRs during slow-wave sleep but distinct from shuffled Non-SWR periods.

**Table S1. Summary of identified slow wave sleep and REM sleep epochs in the 16 analyzed sessions**

| Session | Slow Wave Sleep | | REM Sleep | |
|---|---|---|---|---|
| | Total Time (s) | Count | Total Time (s) | Count |
| 1 | 120 | 6 | 0 | 0 |
| 2 | 150 | 18 | 0 | 0 |
| 3 | 0 | 0 | 0 | 0 |
| 4 | 408 | 39 | 0 | 0 |
| 5 | 44 | 4 | 0 | 0 |
| 6 | 60 | 2 | 0 | 0 |
| 7 | 614 | 20 | 24 | 2 |
| 8 | 178 | 7 | 0 | 0 |
| 9 | 94 | 2 | 0 | 0 |
| 10 | 84 | 11 | 0 | 0 |
| 11 | 296 | 29 | 0 | 0 |
| 12 | 0 | 0 | 0 | 0 |
| 13 | 1024 | 153 | 42 | 2 |
| 14 | 672 | 166 | 250 | 20 |
| 15 | 844 | 19 | 0 | 0 |
| 16 | 286 | 32 | 10 | 1 |

**Table S2. Summary of sorted single-units (i.e., putative hippocampal cells) and decoding accuracy of spiking activity in the 16 analyzed sessions.** Decoding error was measured as the difference between the decoded position and the animal's actual position, using 250 ms decoding windows for analysis.

| Session | Putative principal unit N | | Putative interneuron unit N | | Decoding error (cm) |
|---|---|---|---|---|---|
| | CA1 | DG/CA3 | CA1 | DG/CA3 | |
| 1 | 15 | 33 | 2 | 9 | 18 |
| 2 | 10 | 22 | 2 | 6 | 24 |
| 3 | 20 | 33 | 7 | 9 | 18 |
| 4 | 38 | 52 | 4 | 9 | 18 |
| 5 | 11 | 41 | 2 | 7 | 24 |
| 6 | 4 | 35 | 4 | 6 | 35 |
| 7 | 16 | 1 | 9 | 0 | 33 |
| 8 | 15 | 6 | 4 | 0 | 31 |
| 9 | 23 | 28 | 3 | 5 | 16 |
| 10 | 26 | 34 | 12 | 5 | 13 |
| 11 | 13 | 50 | 3 | 13 | 21 |
| 12 | 7 | 28 | 2 | 19 | 29 |
| 13 | 10 | 29 | 3 | 25 | 18 |
| 14 | 5 | 27 | 2 | 8 | 23 |
| 15 | 4 | 34 | 1 | 6 | 32 |
| 16 | 22 | 31 | 2 | 9 | 19 |
